# A unified network systems approach uncovers a core novel program underlying T follicular helper cell differentiation

**DOI:** 10.1101/2025.08.19.670906

**Authors:** Alisa A. Omelchenko, Syed A. Rahman, Vinayak V. Viswanadham, Grace J. Yuen, Perla M. Del Rio Estrada, Valentino D’Onofrio, Yijia Chen, Na Sun, Hamid Mattoo, Chinmay G. Varma, Gonzalo Salgado, Maribel S. Nava, Lady C. Ruiz, Dafne D. Rivera, Santiago A. Rios, Sudhir P. Kasturi, Susan P. Ribeiro, Mark J. Shlomchik, Amanda C. Poholek, Shiv S. Pillai, Rebecca A. Elsner, Vinay S. Mahajan, Jishnu Das

**Affiliations:** Center for Systems Immunology, Departments of Immunology and Computational & Systems Biology, University of Pittsburgh School of Medicine, Pittsburgh, PA, USA; The Joint CMU-Pitt PhD program in Computational Biology, School of Medicine, University of Pittsburgh, PA, USA; Ragon Institute of MGH, MIT and Harvard, Cambridge, MA; Department of Biomedical Informatics, Harvard Medical School, Boston, MA; Pathology Advanced Translational Research Unit (PATRU), Department of Pathology and Laboratory Medicine, Emory University School of Medicine, GA, USA; Center for Vaccinology, Ghent University and Ghent University Hospital; Integrative Systems Biology PhD Program, School of Medicine, University of Pittsburgh, Pittsburgh, PA, USA; Department of Pathology, Brigham and Women’s Hospital, Boston, MA; Centro de Investigacion en Enfermedades Infecciosas, Mexico City, Mexico; Department of Pathology and Laboratory Medicine, Emory University School of Medicine, GA, USA; Department of Immunology, University of Pittsburgh School of Medicine, Pittsburgh, PA, USA

## Abstract

T follicular helper (Tfh) cells are central to the adaptive immune response and exhibit remarkable functional diversity and plasticity. The complex nature of Tfh cell populations, inconsistent findings across experimental systems and potential differences across species have fueled ongoing debate regarding core regulatory pathways that govern Tfh differentiation. Many studies have experimentally investigated individual proteins and circuits involved in Tfh differentiation in limited contexts, each providing only a partial understanding of the process. To address this, we adopted a novel multi-scale network systems approach that incorporates both regulatory and protein-protein interactions. Our approach integrates diverse data types, captures regulation across multiple levels of immune system organization, and recapitulates known drivers. Further, we discover a core Tfh gene set that is conserved across tissue types and disease contexts, and is consistent across data modalities - bulk, single-cell and spatial. While components of this set have been individually reported, a novel aspect of our work lies in the discovery, characterization, and connectivity of this core signature using a single unbiased approach. Using this method, we also uncover a novel function of IL-12, a molecule with reported conflicting functions, in the regulation of Tfh differentiation. Notably, we find that, in both humans and mice, IL-12 is permissive for the differentiation of Tfh precursors, but blocks subsequent differentiation into GC Tfh cells. Overall, this work elucidates novel networks with unexplored roles in governing Tfh cell differentiation across species and tissues, paving the way for novel -therapeutic interventions.

## Introduction

CD4+ T helper cells are central to the adaptive immune response and exhibit remarkable functional diversity and plasticity^1^. The dynamic nature of T helper cell identity is shaped by complex epigenetic and transcriptional networks and recognized as a critical determinant of immune outcomes^2^. However, understanding the mechanisms governing lineage commitment and plasticity remains a fundamental question in immunology^3–5^. Among T helper subsets, T follicular helper (Tfh) cells represent a particularly complex and heterogeneous population^6^. Tfh cells are essential for supporting B cell maturation within germinal centers (GCs), driving somatic hypermutation, affinity maturation, and long-lived antibody responses^7^. They are indispensable for protective humoral immunity to infection. Dysregulated Tfh cell responses have also been implicated in autoimmunity and cancer ^8–10^.

Despite their importance, inconsistent findings and reported differences across species and experimental systems have fueled ongoing debates regarding the core regulatory pathways that govern Tfh differentiation^11,12^. For example, relying heavily on in vitro culture systems, there have been reports of the necessity of TGF-β and activin A for human, but not murine, Tfh differentiation^12,13^. There has also been a long standing debate on the role of IL-12 in this process when comparing in vitro human data to murine in vivo data across various infectious models^14–19^. These inconsistencies are partially due to the lack of in vitro conditions that produce cells representative of Tfh isolated from lymphoid organs^11^, but further complicated by heterogeneity among Tfh populations that is still poorly defined^20,21^. Part of our gaps in understanding stem from the fact that many studies have experimentally investigated individual proteins and circuits necessary for Tfh differentiation in specific contexts, each providing only a limited window into the process^22–26^. In the absence of a unified systems approach to analyzing these disparate datasets, it currently remains unclear how much of the observed inconsistencies arise from technical artifacts versus fundamental biological differences (if any) between human and murine Tfh differentiation.

To address this issue, we adopted a novel multi-scale network systems approach that incorporates both regulatory and physical interactions^27^. Our approach integrates diverse data types and captures regulation across multiple levels of immune system organization^28,29^. At the transcriptional level, gene regulatory networks (GRNs) provide insight into how genes coordinate and execute immune responses, leading to the discovery of novel regulatory components^30,31^. In addition, protein-protein interactions are also critical regulators of differentiation. These interactions transmit extracellular contextual cues, such as local inflammatory cytokines or cell-cell interactions, through physical associations of ligands with their receptors. Thus, simultaneously incorporating the protein-protein interaction (PPI) network provides orthogonal insights^32^. The PPI network, built from experimentally validated physical interactions, enables the identification of important protein groups with a specific connectivity structure based on their physical associations (i.e., modules). We use network approaches to systematically elucidate circuits underlying Tfh differentiation by integrating data from a set of comprehensive ‘omic profiles. We then interrogate the identified circuits across data types, disease contexts and species to evaluate the generalizability of the identified circuits as well as its conservation across species.

Specifically, we first demonstrated that our novel multi-scale network-based approach can integrate different data modalities from a single set of experiments and identify known regulators of Tfh differentiation that were assembled from many studies. Applying this approach to more diverse datasets, we then discovered that there is indeed a core Tfh gene set which is conserved across tissue types and disease contexts, and consistent across data modalities (bulk, single-cell and spatial). While several components of this set have been individually reported, a unique aspect of our work lies in the discovery and characterization of this core signature using an unbiased approach. More importantly, we found that this core set is conserved across humans and mice, addressing a fundamental open question in the field, as different systems (in-vitro, ex-vivo and in-vivo) have often led to different components being discovered. Further, the core Tfh gene set is functionally linked to specific immune signaling pathways. In this context, we uncovered a novel function of IL-12 in the regulation of Tfh heterogeneity. IL-12 specifically blocks the differentiation of Tfh precursors into GC Tfh cells, in both humans and mice, consistent with recent studies indicating that they correspond to transcriptionally distinct cell states within the Tfh differentiation trajectory^33,34^. Overall, we uncover networks with unexplored roles in governing Tfh cell differentiation across species and tissues, paving the way for novel -therapeutic interventions.

## Results

### An unbiased network systems approach empowers discovery of modules underlying Tfh differentiation

We characterized the epigenomic and transcriptomic landscapes of Tfh cells isolated from adult healthy human tonsils at distinct stages of differentiation. We profiled chromatin accessibility with ATAC-Seq^35,36^, gene expression with RNAseq, and histone H3 modifications with CUT&RUN^37^ using antibodies against H3K4me3, H3K4me1 and H3K27Ac to map promoters, enhancers and their overall chromatin activation respectively. Each of these analyses was performed at four stages of Tfh cell differentiation - Naive (CD45RA+CXCR5-PD1-), Early Extra-GC Tfh/Pre-Tfh (CD45RA-CXCR5+PD1-), Late Extra-GC Tfh/Pro-Tfh (CD45RA-CXCR5+PD1+), and GC Tfh cells (CD45RA-CXCR5++PD1++) (Figure 1A). The CD45RA-CXCR5+PD1- and CD45RA-CXCR5+PD1+ stages express intermediate levels of CXCR5, PD1 and BCL6 compared to naive and mature Tfh cells, suggesting they represent intermediate stages in the Tfh differentiation pathway^38^. This resource represents a global epigenomic roadmap of human Tfh cell differentiation states (Figure 1B). Low-dimensional representations of data from each chromatin assay revealed distinct stage-specific profiles across the four cell subsets that positioned them along a trajectory from the least to the most differentiated state, reflecting the stepwise activation of regulatory programs during Tfh cell differentiation (Extended Figure 1A, 1B). Here, we adopted a novel multi-scale network approach to integrate the different omic datasets as each individual dataset typically captures only a partial view of the complex and interconnected immune networks underlying immune responses (Figure 1C). This fragmented perspective can obscure underlying biological mechanisms and fuel scientific disagreements, as researchers interpret different slices of the data landscape. Network and systems based approaches help overcome this challenge by integrating diverse modalities to generate multiscale, unbiased, and interpretable views of immune responses ^28,29,39^. Further, although bulk datasets lack cellular resolution^40^, when paired with biological networks they can support discovery by identifying functional modules and key regulators (Figure 1D). Incorporation of single cell and spatial information further empowers refinement of the core components. It also allows for meaningful inference and identification of targets which can be experimentally validated using in vivo murine models (Figure 1D).

**Figure 1:**
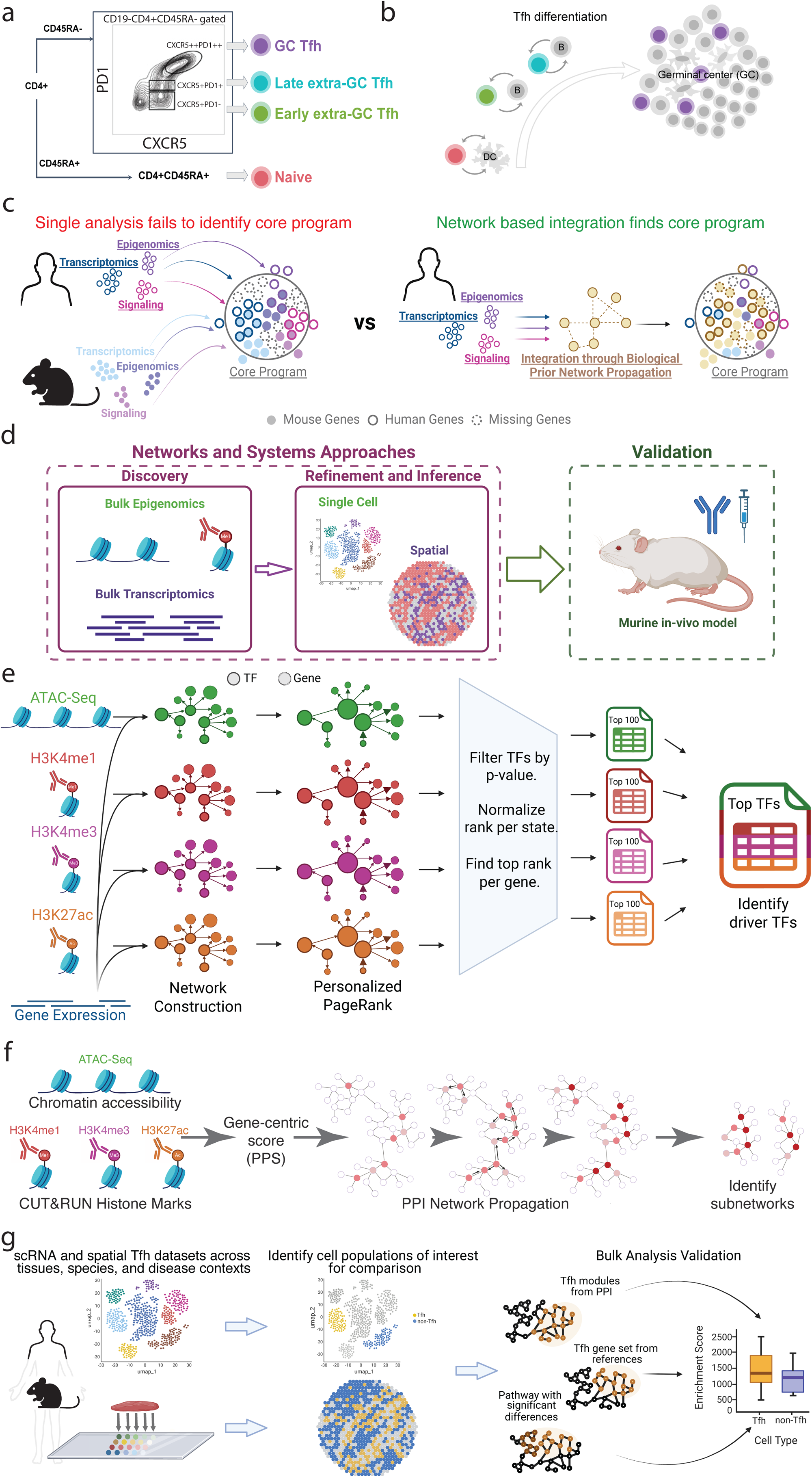
A. Flow cytometry gating strategy to identify Tfh cells and their differentiation state from the 600 million human tonsil cells. B. Tfh cell differentiation trajectory of the 4 states identified through flow cytometry. C. Schematic displaying multi-omic datasets each providing a specific view of a system and how using networks as biological priors can integrate the views and identify a core program across different conditions such as species. D. Overview showing how novel targets were discovered using bulk datasets and refined using single cell and spatial data. Targets were then validated in a murine in-vivo model. E. Method for identification of top driver TFs through integration of the epigenomic/ transcriptomic datasets with Personalized PageRank network propagation and filtering the output ranks. F. Protein-protein interaction network propagation using the gene centric peak proximity score (PPS) to integrate epigenomic datasets and identify significant subnetworks. G. Validation of bulk data results with single cell RNA (scRNA) datasets across contexts like disease and species through single cell gene set enrichment analysis (scGSEA).

Using these datasets, we first identified prioritized TFs underlying Tfh differentiation using a personalized PageRank approach that leverages gene regulatory network (GRN) architecture, where the network is identified by combining the transcriptomic and epigenomic data (Figure 1E). However, GRNs capture only one dimension of the regulatory landscape and physical protein interactions, such as receptor-ligand associations, also play a critical role in cellular differentiation. We therefore also included in our integrative framework a protein-protein interaction (PPI) network-centric approach that revealed additional components of the core regulatory program not captured by GRN-based approaches alone (Figure 1F). For the PPI-based approach, we integrated signals across a range of datasets using a novel network propagation technique that integrates diverse epigenomic signals (that are potentially individually noisy) into a robust set of modules. We first identified chromatin peaks, i.e. regions enriched for accessible chromatin or histone marks, that significantly varied across all four stages of Tfh differentiation (Extended Figure 1B) and applied a gene-centric approach to integrate these epigenetic signals at the level of individual genes (Extended Figure 1C).

Our combined strategy thus prioritizes genes based on both their regulatory impact and their physical interaction context. To validate and refine the core transcriptional programs identified from bulk population data, we analyzed single-cell RNA-seq (scRNA-seq) and spatial datasets spanning diverse tissues, species, and disease contexts (Figure 1G). Although scRNA-seq provides cellular resolution, it suffers from technical noise and dropout, whereas data from cell populations offers higher signal fidelity, enabling robust identification of core regulatory modules via network propagation. This integrative approach helped us to cross-validate findings from bulk population data across species and platforms, including single-cell and spatial datasets, and to prioritize conserved regulatory genes. We then benchmarked our gene sets against established Tfh signatures to ensure specificity and biological relevance.

### Network-based integration of epigenomic and transcriptomic data uncovers core processes driving Tfh cell differentiation

We first applied a framework leveraging cell-state-specific GRNs to identify key transcription factors (TFs) driving Tfh differentiation. Specifically, we used a Personalized PageRank approach to construct GRNs from epigenomic data^41^, and then implicate TFs that distinguish between the different stages of Tfh differentiation. From each epigenomic dataset, we selected the top 100 TFs based on their highest rank across all stages, yielding a combined set of 152 unique TFs from an initial pool of ∼1,200 candidates (Figure 2A, 2B, Methods). Notably, the top-ranking TFs showed only weak correlation with gene expression changes (Figure 2A, y-axis), highlighting a critical limitation of relying solely on differential gene expression to prioritize TFs. This disconnect between differential gene expression and regulatory significance may be particularly relevant to the Tfh context, where key regulators of Tfh differentiation including BCL6 are subject to tight post-transcriptional and translational control^42^. Indeed, only a small number of genes encoding TFs with high mRNA fold change (log_2_FC) of expression were ranked highly by Taiji and passed standard significance thresholds.

**Figure 2:**
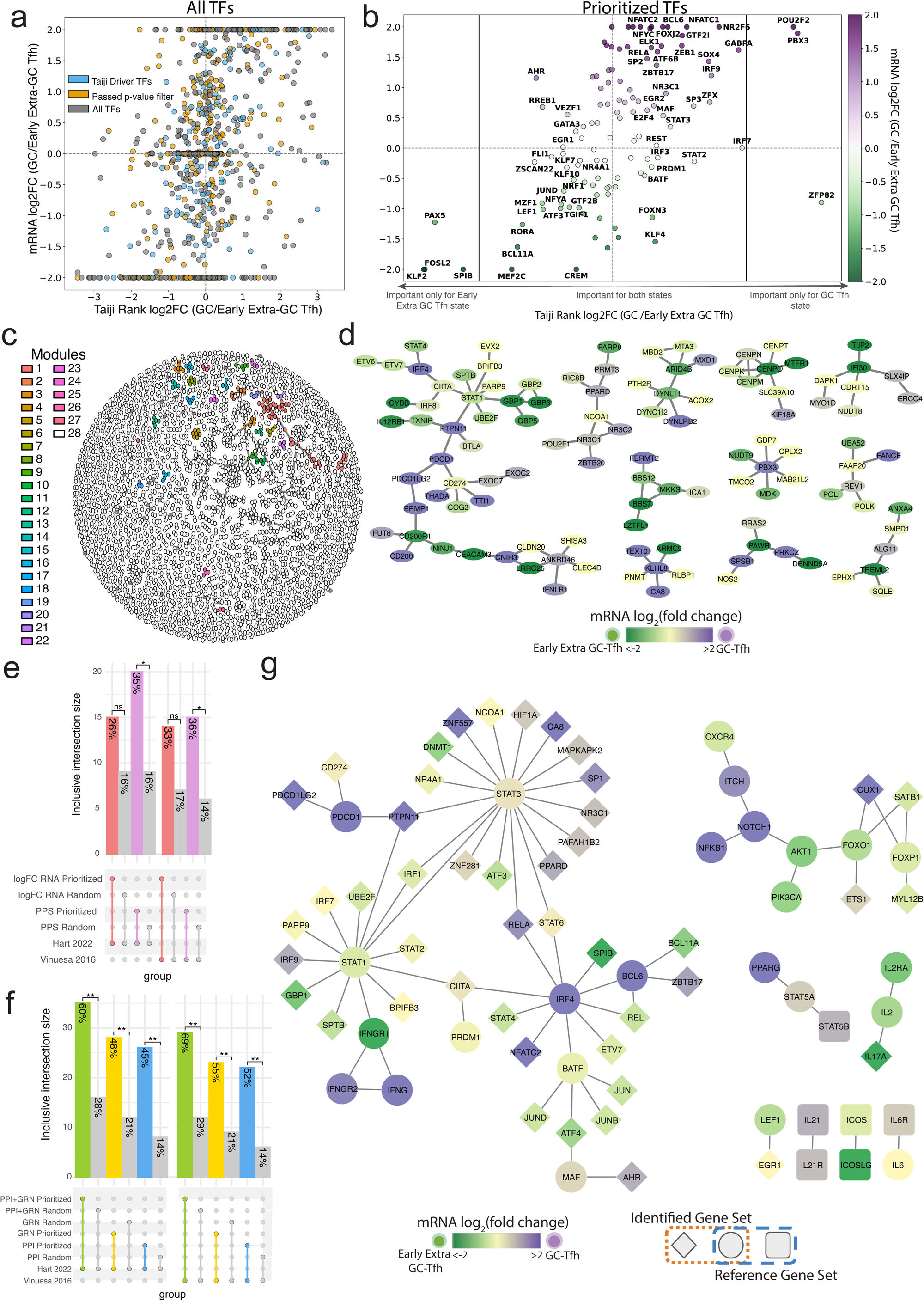
A. The spread of the 1165 TFs analyzed by Taiji given by the mRNA log_2_FC between the GC and Early Extra GC state (y-axis) and the Taiji Rank log_2_FC between the 2 states (x-axis) with all TFs (grey), those which passed the p-value threshold of <0.01 (orange), and the chosen 152 driver TF set (blue). B. A focused scatter plot on the driver TF set without ZNFs analyzed by Taiji given by the mRNA log_2_FC between the GC and Early Extra GC state (y-axis) and the Taiji Rank log_2_FC between the 2 states (x-axis) and colored by the mRNA log_2_FC of the GC and Early Extra GC Tfh where higher expression in GC is shown in purple, and higher in Early Extra GC Tfh is shown in Green. C. Visualization of 2000 random edges (module 28) and the subnetworks identified through the PPI network propagation (modules 1-27) showing specificity of the selected modules. D. Largest connected modules (>6 genes), colored by the mRNA log2FC as described in B. E. Overlap of size matched gene sets of log_2_FC RNA and top PPS with their first-degree interactors with 2 reference Tfh sets. Size matched sets (184 genes) of the top log_2_FC RNA scores or PPS were identified and first-degree interactors were added from the network. 1000 permutations of a random size matched sets were also compared for each analysis. Fisher’s exact test performed with average random overlap, P-value threshold <0.05. F. PPI, GRN, and PPI + GRN prioritized gene sets plus direct interactors overlap with 2 Tfh reference gene sets. 1000 permutations of a random size matched sets were also compared for each analysis. Fisher’s exact test performed with average random overlap, P-value threshold <0.05. G. Overlay of the PPI+GRN prioritized set with direct interactors on the Vinuesa 2016 reference network. Shared genes (circle), analysis only discovered interactors (diamond), and reference only genes (square) are colored by mRNA log_2_FC as previously described.

We examined the stage-specific relevance of key TFs by looking at the difference in rank across the early extra-GC and GC Tfh states (Figure 2B, left to right on the X axis: moves from only relevant for early extra-GC state to relevant for both states to relevant for GC Tfh state only). KLF2 and FOSL2 were selectively ranked as only important for the early extra-GC Tfh state. This is consistent with prior work that demonstrates that KLF2 is widely expressed on naive mature CD4+ T-cells and repressed in GC Tfh cells ^43,44^. Similarly, miR-155 mediated repression of FOSL2 has been shown to be important for Tfh cell fate commitment ^45,46^. In contrast, we find that POU2F2 and PBX3 were important only in the GC-Tfh state. The role of POU2F2 (OCT2) in promoting germinal centers as well as GC B-cells has been extensively studied^47,48^. Limited studies have shown POU2F2 to be involved in Tfh differentiation through transactivation of the BCL6 and BTLA promoters^49,50^. This analysis suggests that POU2F2 may play a more profound role specifically in GC Tfh differentiation, as opposed to other Tfh states. We additionally identified a putative novel factor, PBX3, which has not been studied in GC Tfh differentiation to our knowledge. Other TFs, such as BCL6, were important in both contexts. Although BCL6 shows a strong expression log₂FC favoring the GC-Tfh state, its ranking underscores its broader relevance as a lineage-defining TF for Tfh cells^51^. PRDM1, a well-known repressor of the Tfh program^51^, exhibits minimal transcript-level changes. Despite this, it was still correctly prioritized as a key regulator of the Tfh program in both states as it is well-known that PRDM1 needs to be turned off (along with BCL6 being turned on in a reciprocal loop) for the formation of GC Tfh^51^. Conventional expression-based analyses would have missed most of these hits, further demonstrating the value of using this regulatory network-informed approach.

### PPI integration identifies submodules in Tfh cell differentiation

Next, to move beyond GRNs, which do not capture physical interactions such as receptor-ligand binding and signaling cascades that also play critical roles in cell differentiation, we integrated a high-confidence human PPI network into our epigenomic analyses. The validity of each edge in our PPI network is supported by multiple independent experimental assays^52,53^. However, the PPI network is context agnostic. We added context-specific data to this network by using each protein’s node weight (score for the gene encoding the protein based on its aggregate PPS computed using all chromatin features)^54^. Stage-specific epigenetic signals for each chromatin feature were mapped to nearby genes and combined into a single “peak proximity score” (PPS). The PPS was calculated as a distance-weighted sum of the number of variable CUT&RUN or ATAC-Seq peaks relative to a transcription start site, with equal contribution from each chromatin feature. The PPS, a gene-centric measure of stage-specific epigenetic activity, exhibited significant correlation with gene expression levels in both Early Extra-GC and GC Tfh stages, but the magnitudes of the Spearman correlation were modest (Extended Figure 1D, E).

This information was then propagated along the network using a random walk with restart approach. There are two key goals achieved by the network propagation approach. First, it identifies high-scoring subnetworks in the network, taking into account additive effects based on functional similarity (with edges in the networks as surrogates for functional similarity) rather than focus on individual high-scoring nodes. Second, due to false negatives (signal not captured) inherent to omic assays, this detection of high-scoring networks occurs after propagating the initial signal to account for initial incompleteness in the data. The discovery of meaningful regulatory modules depends upon both the input scores and the structure of the interaction network. To ensure that only robust and biologically grounded modules were retained, we selected statistically significant subnetworks using stringent permutation testing. Modules identified using randomized gene scores or shuffled networks were excluded ensuring that the selected subnetworks reflected true biological signals. The prioritized modules were consistently identified across five independent network propagation replicate runs. By leveraging the topology of the PPI network as prior knowledge that encodes biological dependencies (functional similarity), this approach allowed us to identify coherent subnetworks that may underlie Tfh cell differentiation.

The resulting 27 modules, comprising 184 genes, map to distinct regions of the interactome and represent <1% of the full network (Figure 2C). We overlaid the mRNA log_2_FC between GC and Early Extra-GC Tfh populations to distinguish which genes are more highly expressed in each differentiation state (Figure 2D, Extended Figure 2A). Most modules span both states, capturing the dynamic regulation across the Tfh trajectory. Notably, the largest connected module includes several canonical Tfh-associated genes such as CD200 and PDCD1, underscoring its biological relevance (Figure 2B). In contrast, some modules exhibit clear state-specificity. For example, a module enriched for IL17 signaling components, including IL17RC, IL17F, IL17RA, IL17A, and IL2 shows comparatively elevated expression in the Early Extra-GC Tfh state (Extended Figure 2A).

Building on the identified subnetworks and ranked transcription factors, we integrated the driver TFs, the module genes, and their direct interactors within the protein network to define a final set of prioritized genes (Methods). To assess the biological relevance of this gene set, we evaluated its overlap with proteins annotated in multiple experimentally validated human Tfh reference datasets^55,56^. Since some of the overlap could be explained by the input transcriptomic and epigenomic signals alone, we generated size-matched gene sets using either top-ranked genes from RNA-seq log₂FC or from epigenomic PPS scores, along with their direct protein interactors (Methods). For all sets, we also generated 1,000 permutations of random size-matched controls to establish a random baseline for expected overlap. We found that the transcriptomic and the epigenomic data alone only weakly recovered biologically relevant genes (Figure 2E, Extended Figure 2B). These results indicate that while gene-centric scores provide useful starting points, they are incomplete on their own.

We found that network-based approaches showed substantially greater overlap with experimentally validated Tfh reference datasets, with enrichment levels between 2- to 4-fold above the random baseline (Figure 2F, Extended Figure 2B). Notably, the combined network approach outperformed individual methods, recovering 60–69% of known Tfh-associated genes compared to 45–55% for individual subnetworks or TF rankings alone. These results highlight the added value of integrating multiple network-derived signals. Together, this demonstrates that network propagation provides a critical framework for identifying higher-order regulatory relationships that are not captured by transcriptomic or epigenomic data alone, and that combining orthogonal network-based strategies further improves the ability to recover biologically meaningful targets.

To understand how the curated genes from literature are connected with each other, higher order information must be provided. Here, we use a protein interaction network to provide the functional relationship of these genes. We visualized the Vinuesa 2016 reference network by identifying how they are connected in the broader PPI interactome. We then overlaid our prioritized Tfh gene set onto this network and included any direct interactors of the reference genes that were also part of the prioritized set (Figure 2G). Several key regulators of Tfh differentiation, such as BCL6, PRDM1, STAT1, STAT3, and PDCD1, were successfully recovered (denoted as circles). In addition, many biologically relevant genes not included in the original reference set (shown as diamonds) served as connectors within the network. For example, PTPN11 (SHP2) links PDCD1 to STAT3 and STAT1, highlighting potential intermediates that bridge well-established signaling nodes^57^. When overlaying the top mRNA logFC identified genes with their direct interactors, the resulting network is disjointed (Extended Figure 2C). While STAT1, STAT3, and IRF4 were identified in the mRNA logFC genes, their connections to each other through interactors such as STAT6, RELA, or CIITA have been lost. Moreover, we can visualize the rank importance and module changes across the differentiation trajectory by also comparing to the Late Extra-GC Tfh state (Extended Figure 2D, 2E). This demonstrates the power of using network propagation to give context to higher order organization across the Tfh differentiation trajectory.

These results also illustrate that our prioritized gene set can capture both known regulators and novel or less-characterized players in Tfh cell biology. However, some well-known Tfh-associated genes, including IL21, CXCR5, and ICOS, were not identified through our approach. Since these are not TFs, these genes would have to be found through the PPI network. We analyzed the shortest PPI paths from the reference set and found that although these genes initially carried strong signals, they were directly connected to interaction hubs. This caused signal dissipation during propagation. For instance, IL21 and IL21R are connected to SMARCA2, which interacts with 132 proteins (Extended Figure 2B). Similarly, CXCR5 is connected to GNAI2, which interacts with 72 proteins and is linked to VIRMA, a node with over 2,700 connections (Extended Figure 2C). ICOS also interacts directly with VIRMA. These findings underscore an important limitation of PPI-based propagation: while the method effectively controls false positives, it may have some false negatives due to data, but not methodological, limitations.

### Single cell spatial and RNAseq datasets align with network prioritized gene sets over curated Tfh reference sets

We hypothesized that if the modules derived from bulk omics datasets are truly reflective of the GC Tfh state, they should consistently exhibit higher enrichment scores in Tfh populations relative to non-Tfh cells across independent single-cell and spatial datasets. Therefore, we examined how well the modules identified earlier from bulk data extrapolate to scRNAseq and spatial datasets spanning diverse tissue types and contexts.

First, we acquired and processed scRNAseq data from human gut samples from 4 tissue locations of people living with HIV (PLWHIV) (Figure 3A). The Tfh cell population was identified through markers and the Azimuth reference datasets (Figure 3B, Methods). Along with Tfh cells, two non-Tfh populations, Central Memory (CM) non-Tfh CD4+ and Th17, were chosen for comparative analyses (Figure 3C). We implemented single cell gene set enrichment (scGSEA) analyses. scGSEA is a statistical method designed to measure the composite effect of a predefined gene set at single-cell resolution^58^. Each prioritized gene set found with the network approaches was evaluated. We also benchmarked against 2 previous reviews summarizing Tfh drivers from a large number of studies (the Vinuesa 2016 and Hart 2022 literature reviews). The signal in each of these sets was evaluated using a combination of a statistical significance and an effect size (Cliffs Δ, a non-parametric effect size measure used to quantify the difference between 2 groups) threshold. When comparing the Tfh and CM non-Tfh CD4+ populations, both the PPI and GRN prioritized gene sets (Cliffs Δ = 0.27, 0.15 respectively, considered moderately large effect sizes for heterogeneous single-cell data) outperformed the Tfh reference sets (Cliffs Δ = -0.1 and 0.05 respectively) (Figure 3D). The evaluation of the Tfh and Th17 populations show comparable effect sizes for all gene sets (Figure 3E). This demonstrates that while curated gene lists can separate the Tfh population from some cell types like Th17, they fail to distinguish Tfh populations from other cell types like CM non-Tfh CD4.

**Figure 3:**
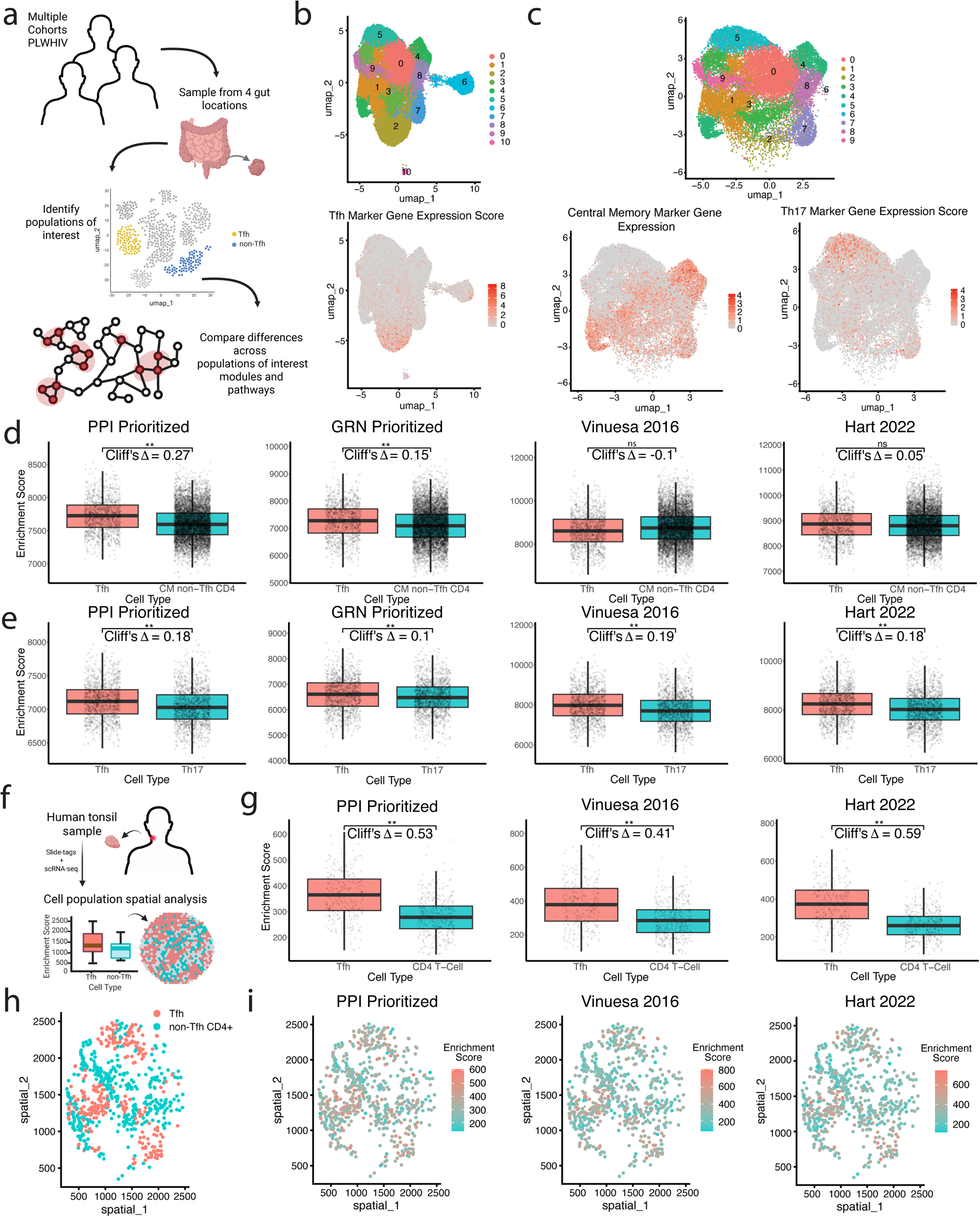
A. Illustration of the refinement analysis using a Tfh and non-Tfh cell population identification and gene set enrichment analysis for modules of interest. Cell populations were extracted from a human scRNA dataset composed of gut samples from people living with HIV (PLWHIV). B. UMAP visualizing scRNA clusters (top) made using shared nearest neighbor (SNN) graph construction and the Louvain algorithm, with parameters k.param= 50 and resolution = 0.7 based on the Harmony-reduced space. Tfh cell identification using Tfh Marker Gene Expression Score. C. Identification of non-Tfh populations from the scRNA clusters (top) after filtering and removing cell populations using the Central Memory (bottom left) and Th17 (bottom right) Marker Gene Expression Score. D. scRNA enrichment for CM non-Tfh CD4 and Tfh cell populations for PPI prioritized, GRN prioritized, Vinuesa 2016 published reference, and the Hart 2022 published reference gene sets (Cliffs Δ = 0.27, 0.15, -0.1, 0.05). Significance given by effect size (Cliffs Δ) ≥ 0.1 and p-value > 0.01. E. scRNA enrichment for Th1, GRN prioritized, Vinuesa 2016 published reference, Hart 2022 published reference gene sets (Cliffs Δ = 0.18, 0.1, 0.19, 0.18). Significance given by effect size (Cliffs Δ) ≥ 0.1 and p-value > 0.01. F. Schematic demonstrating the analysis of the human tonsil spatial dataset. G. scRNA enrichment for non-Tfh CD4+ and Tfh cell populations for PPI prioritized, Vinuesa 2016 published reference, and the Hart 2022 published reference gene sets (Cliffs Δ = 0.53, 0.41, 0.59). Significance given by effect size (Cliffs Δ) ≥ 0.1 and p-value > 0.01. H. 2D visualization of the spatial coordinates of non-Tfh CD4+ and Tfh cells colored by cell type. I. 2D visualization of the spatial coordinates of non-Tfh CD4+ and Tfh cells colored by enrichment score for the PPI prioritized, Vinuesa 2016 published reference, and Hart 2022 published reference gene sets.

Next, we examined how well the identified modules defined Tfh cells profiled from healthy human tonsils using spatial transcriptomics (Slide-tag)^59^. As earlier, we defined Tfh and non-Tfh CD4 T-cells using marker genes. The identified modules as well as the reference literature-curated gene sets (Vinuesa 2016 and Hart 2022) were then evaluated in terms of their ability to stratify the Tfh and non Tfh CD4s (Figure 3F). All three sets provided excellent stratification in this case (Figure 3G). Further, we also visualized the spatial location of the cells of interest colored by their cell type (Figure 3H). All gene sets captured relevant spatial microniches of the Tfh cells based on their corresponding enrichment scores (Figure 3I). The ability of the bulk-derived gene sets to distinguish the Tfh population from other cell types when equally or better compared to curated gene sets shows the power of these network approaches. Hence, high-quality bulk datasets in tandem with network approaches can enable capturing complex regulatory processes in the immune system.

### Prioritized gene sets are consistent across species

While in vitro culture systems are necessary for the study of human Tfh differentiation, murine systems are necessary to experimentally interrogate this process in vivo. Current approaches have yielded different results across in-vitro, ex-vivo, and in-vivo systems. This has led to a fundamental question as to whether there are indeed key differences across human and murine Tfh differentiation, or if the observed differences are simply a reflection of artifacts across the different systems. To address this, we compared our human-derived set of core Tfh modules to a manually curated murine Tfh reference set derived from literature^23^. Intriguingly, the identified genes exhibited a comparable degree of overlap with the mouse reference set as it did with human datasets (Figure 2F, Figure 4B). This suggests that there is strong evolutionary conservation between human and murine Tfh differentiation processes. To further test this, we employed a scRNAseq dataset from a murine model of allergic asthma (where mice were continually exposed to intranasal house dust mite)^60^. This in vivo model elicits a strong type 2 immune response and induces germinal center formation in the draining lymph nodes, providing a physiologically relevant setting to assess Tfh identity. scGSEA analyses revealed that our prioritized gene set performed as well as the murine Tfh reference in distinguishing Tfh cells from resting T cells within this allergic microenvironment (Figure 4C). Interestingly, the prioritized gene sets were about twice as good at separating these 2 cell states when compared to the 2016 human Tfh reference gene set, further demonstrating that it captures known and novel core network modules across species and tissues (Figure 4D). Together these results suggest that while conventional methods may fail to reconcile differences between different systems, multimodal approaches can enable the identification of gene modules that are truly conserved across species, tissues, and disease contexts. The consistency of these findings across bulk, single-cell, and spatial transcriptomic datasets supports their robustness and highlights a core gene program that transcends individual experimental platforms.

**Figure 4:**
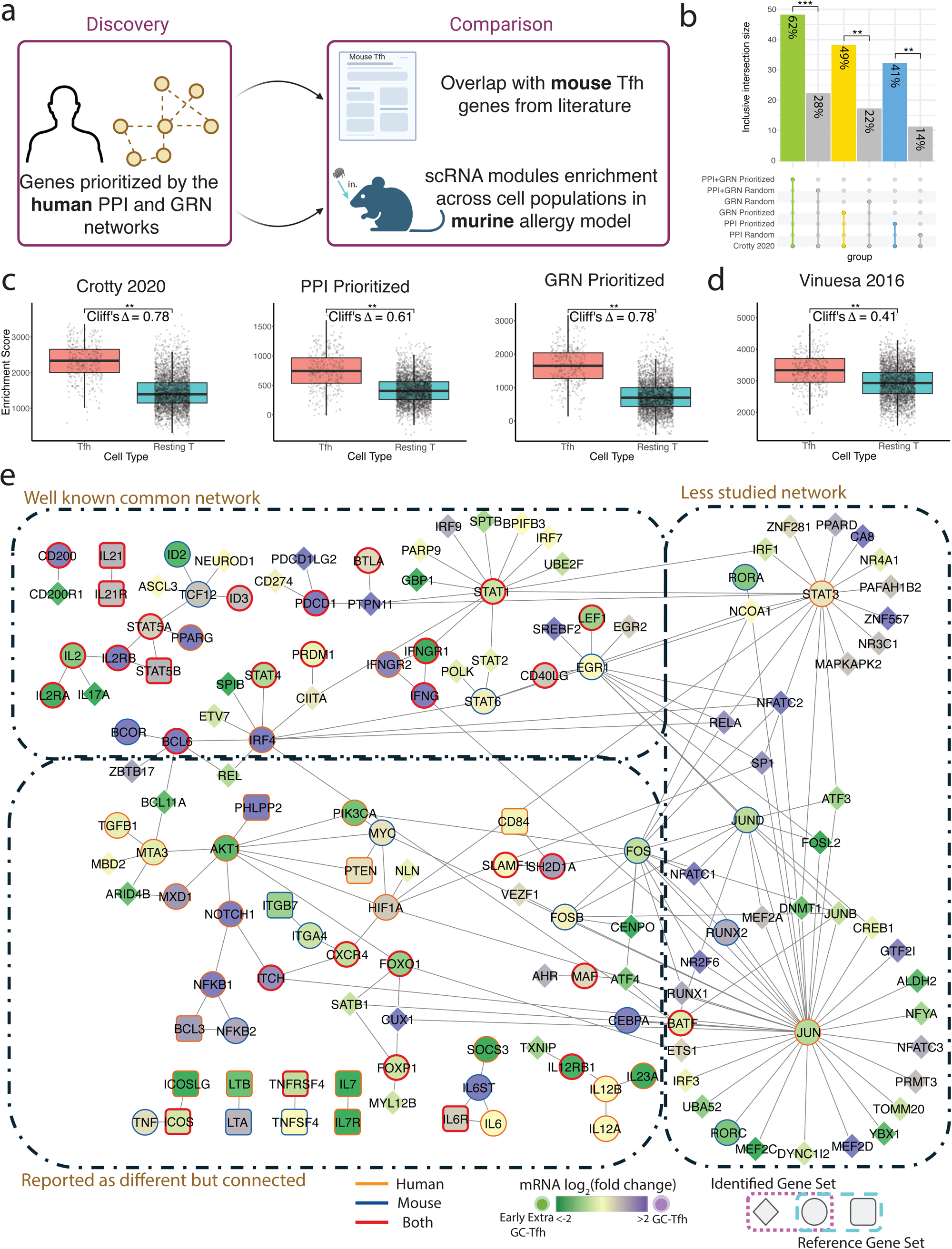
A. Schematic of the experimental set up to identify commonalities between the human derived gene set to mouse networks across data modalities and allergy. B. PPI, GRN, and PPI + GRN prioritized gene sets plus direct interactors overlap with a mouse Tfh reference gene set. 1000 permutations of a random size matched sets were also compared for each analysis. P-value <0.001 and <0.01 respectively. C. scGSEA enrichment for Resting CD4 T-cell and Tfh cell populations for the 2020 murine published reference, PPI prioritized, GRN prioritized gene sets (Cliffs Δ = 0.78, 0.61, 0.78). Significance given by effect size (Cliffs Δ) ≥ 0.1 and p-value > 0.01. D. scGSEA enrichment for Resting CD4 T-cell and Tfh cell populations for the 2016 human published reference (Cliffs Δ = 0.41). Significance given by effect size (Cliffs Δ) ≥ 0.1 and p-value > 0.01. E. Visualization of the identified core Tfh program across mouse and human datasets. Genes from the 2020, 2016, and 2022 references were pooled and the edges between the genes were provided by the PPI network. Shared mouse and human reference genes are outlined red while mouse reference only or human only were outlined in blue and orange respectively. The identified gene set was overlaid with this network and direct interactors were kept. Shared genes (circle), analysis only discovered interactors (diamond), and reference only genes (square) are covered by mRNA log_2_FC as previously described. The network was partitioned into 3 sections: well-known common network, less studied network, and reported as different but connected.

Finally, to visualize the broader architecture of the Tfh cell differentiation landscape, we integrated the murine and human reference gene sets onto the PPI network. Genes present in the human reference were outlined in orange, those in the mouse reference in blue, and those shared between both in red. We then overlaid our prioritized gene set along with any direct interactors of the reference genes included in the network. This integrated network naturally resolves into three conceptual modules (Figure 4E). The first is a well-characterized core set of canonical Tfh regulators (including TFs, ligands and receptors) such as PDCD1, STAT1, CD40LG, BCL6, CD200, IL21, STAT4, and PRDM1. The second is a less-explored region composed of interactors, many linked to transcription factors such as STAT3 and JUN, that have not been highlighted previously as Tfh-associated. Lastly, we identify a “reported as different between species but connected” region which is composed of genes identified in only one species’ reference set but shown by our network analysis to be physically and functionally connected in the integrated network. These findings underscore the value of a network-based approach in unifying seemingly divergent species-specific results and highlighting evolutionarily conserved components of the Tfh regulatory circuit.

### Identified modules correspond to specific immune signaling pathways

Next, we evaluated how our bulk derived prioritized genes related to known biological pathways^61,62^. This provides an orthogonal way to contextualize our findings as the modules are discovered based on the structures of the underlying gene regulatory and protein interaction networks, but do not take into account known pathways or gene sets. We began by assessing overlaps of the modules with the Hallmark gene sets that represent well-defined biological states and processes^63^. To determine significance of overlaps, the observed intersections to those expected from size-matched random sets of network proteins were compared. This provides a rigorous null model (baseline) for expected overlap by chance.

The results were visualized using a volcano plot. Each point represents a specific pathway, with the effect size (log odds ratio) plotted on the X-axis and significance (–log_10_Q) on the Y-axis, reflecting both the strength and statistical support of each enrichment (Methods). Three of 50 pathways passed the thresholds (-log_10_Q > 0.7 and log odds ratio > 0.58) - IL-6/JAK/STAT3 signaling, MYC signaling and TNFα signaling via NF-κB (Figure 5A).

**Figure 5:**
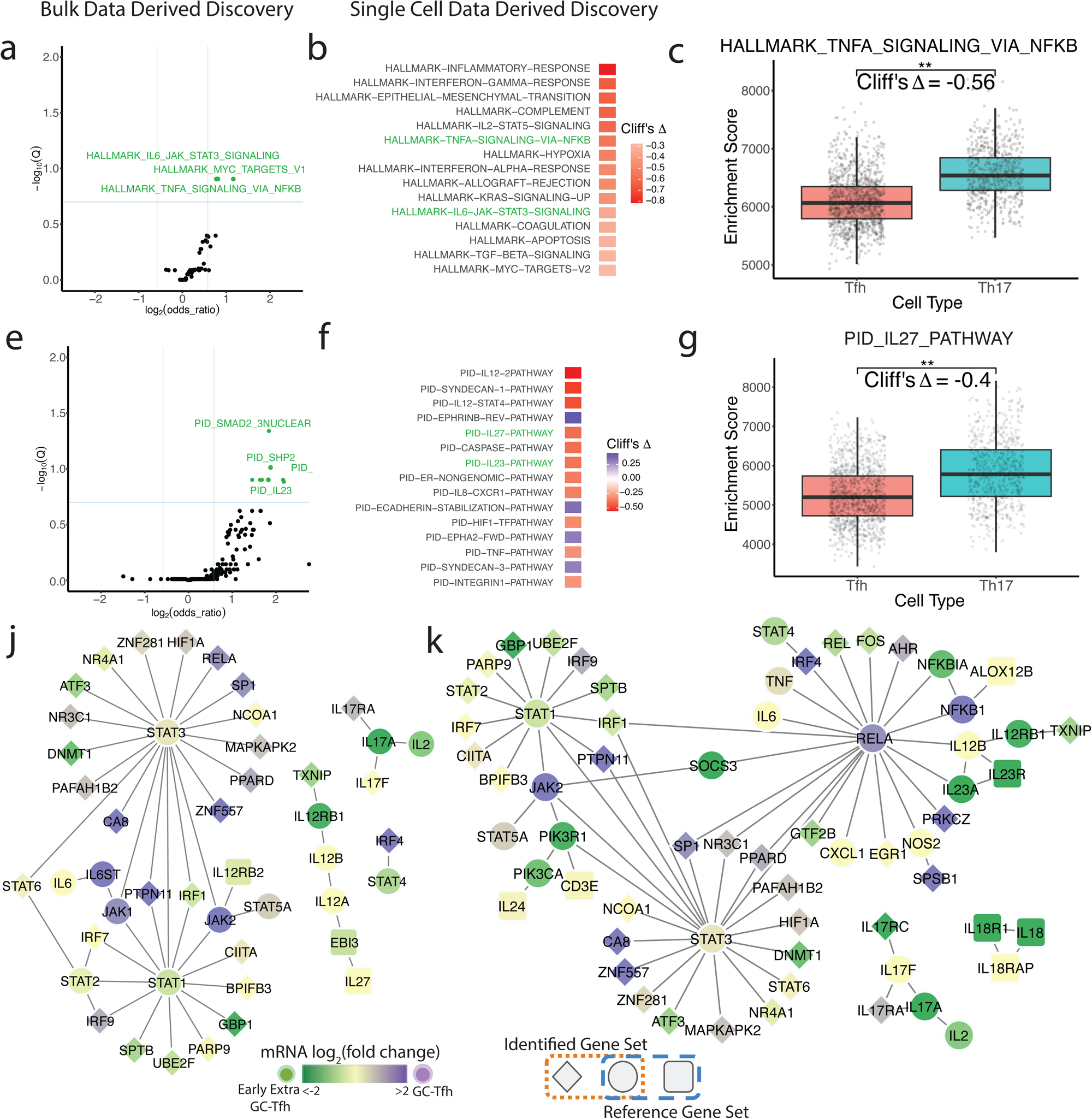
A. Volcano plot visualizing effect sizes and significance of functional overlaps between the Tfh identified gene set and Hallmark pathways. Significant pathways colored in blue (- log(Q-value) > 0.7, log2(odds_ratio) > 0.58). Adjusted p-value (FDR) < 0.2, and 1.5-fold enrichment. B. Human scRNA top 15 Hallmark pathways enriched for the Tfh identified gene set given by the top effect size between the Tfh and Th17 cell population enrichment scores. Pathways that were also identified in the volcano plot are highlighted in blue. C. scGSEA enrichment of the TNFA-SIGNALING-VIA-NFKB hallmark pathway for Th17 and Tfh cell populations (Cliffs Δ = -0.56, p-value < 0.01). D. Volcano plot visualizing effect sizes and significance of functional overlaps between the Tfh identified gene set and PID pathways. Significant pathways colored in blue. Significant pathways colored in blue (-log(Q-value) > 0.7, log2(odds_ratio) > 0.58). Adjusted p-value (FDR) < 0.2, and 1.5-fold enrichment. E. Human scRNA top 15 PID pathways enriched for the Tfh identified gene set given by the top effect size between the Tfh and Th17 cell population enrichment scores. F. scGSEA enrichment of the IL27 PID pathway for Th17 and Tfh cell populations (Cliffs Δ =-0.40, p-value < 0.01). G. PID IL-27 pathway PPI network visualization with the prioritized Tfh gene set overlaid. Shared genes (circle), analysis only discovered interactors (diamond), and reference only genes (square) are colored by mRNA log_2_FC as previously described. H. PID IL-23 pathway PPI network visualization with the prioritized Tfh gene set overlaid. Shape and color as previously described.

We repeated the bulk and scRNA pathway enrichment analysis using the KEGG database ^64^. From the bulk data, we also observed the functional overlap with IL6-STAT signaling, demonstrating that our analyses robustly capture the same IL6 pathway as defined in two different databases (Extended Data Figure 3B). Additionally, significant functional overlap with the KEGG cytokine receptor signaling pathway was identified in both the bulk and scRNA analysis (Extended Data Figure 3B,3C).

To further validate our findings, we used the human PLWHIV scRNA-seq dataset to identify Hallmark pathway differences between the Tfh and Th17 cell populations. We used the top Cliff’s Δ to identify the 15 most different pathways across these cell types (Figure 5B). IL-6/JAK/STAT3 and TNFα signaling via NF-κB were also within the top 15. Interestingly, recent findings have linked aberrant extra-follicular TNFα accumulation in COVID-19 to loss of Tfh cells and germinal centers ^65^. We also note the decrease of TNFα in PLWHIV Tfh populations compared to Th17 populations (Figure 5C). Moreover, we visualized our prioritized gene set and direct interactors with the TNFα signaling via NF-κB hallmark pathway (Extended Figure 3A).

Finally, we also evaluated overlaps of the Tfh modules with the PID pathways, which represent human signaling and regulatory processes^66^. The bulk analysis identified a few pathways including SMAD2, which is a well-known signaling pathway in Tfh differentiation^12,13^ (Figure 5D). Additionally, we see enrichment for the IL-27 and IL-23 pathways, which have been less studied in the context of Tfh. IL-12, IL-23 and IL-27 belong to the same heterodimeric family^67^ and there has been some evidence that they regulate Tfh differentiation ^14,68,69^. Since IL-23 is well known to enhance Th17 differentiation, it is expected that the IL-23 pathway would be relatively lower. We visualized our prioritized gene set and direct interactors with the IL-27 (Figure 5G) and IL-23 (Figure 5H) PID pathways. As shown, there is a lot of overlap between the genes in these 2 pathways including STAT1, STAT3, and IL12 related genes. We also observed higher expression levels of genes encoding this cytokine family in the Early Extra GC Tfh state.

### IL-12 fine tunes Tfh differentiation

We noted that the IL-12 family cytokines were consistently identified throughout our unbiased analyses as top hits. These were among the top PID pathways from the scRNA pathway enrichment analysis (Figure 5E-J) and were predicted to be important regulators when comparing early Extra-GC Tfh and GC Tfh (Figure 2D, 2G). However, while the analysis indicates the IL-12 pathway is highly important, expression of key signaling proteins STAT4 and IL12RB1 are downregulated in GC Tfh as compared to Extra-GC Tfh (Figure 2G). Several in vitro studies of human Tfh differentiation suggest that IL-12 enhances Tfh differentiation, but these Tfh are in fact CXCR5^low^ PD-1^low^ and therefore may be more representative of an early Extra-GC Tfh stage. Combining these observations, we hypothesized that IL-12 may play a stage-specific role in Tfh differentiation, enhancing early development towards the Tfh fate but suppressing full differentiation to GC Tfh.

To test this experimentally, we turned to a murine model of infection with the bacteria *Salmonella enterica* serovar Typhimurium (STm), which elicits a strong IL-12 response that suppresses GCs and Tfh formation^19,70^. In this system, the IL-12 suppression mechanism is dominant, meaning that it also suppresses the GC response to a concurrently administered but unrelated antigen, which we utilize to track the effect of STm infection on the GC response. In this case we chose nitrophenol haptenated chicken gamma globulin (NP-CGG) for the unrelated antigen. In this system, only infection with live STm, not heat killed (HK) bacteria, is able to suppress GC responses^19,70^.

To examine the stage-specific effects of IL-12 on Tfh differentiation, we administered IL-12-neutralizing antibodies beginning one day prior to, or 3 days after, STm/NP-CGG immunization in this experimental scheme (Figure 6C-E, orange). Blocking IL-12 starting prior to infection ensures T cells neither receive IL-12 at priming nor throughout GC formation. At day 3 T cells have been primed and pre-Tfh have moved to the T/B border to interact with pre-GC B cells, but the GC structure has yet to form and GC Tfh differentiation has yet to occur^33,71,72^. We then examined the effect of either early or day 3-initiated IL-12 blocking on total Tfh (CXCR5^high^ PD-1^high^), and GC Tfh or non-GC Tfh, which are CD90^low^ or CD90^high^, respectively (Figure 6D, Extended Figure 5A)^33^ at 12 days after infection/immunization.

**Figure 6:**
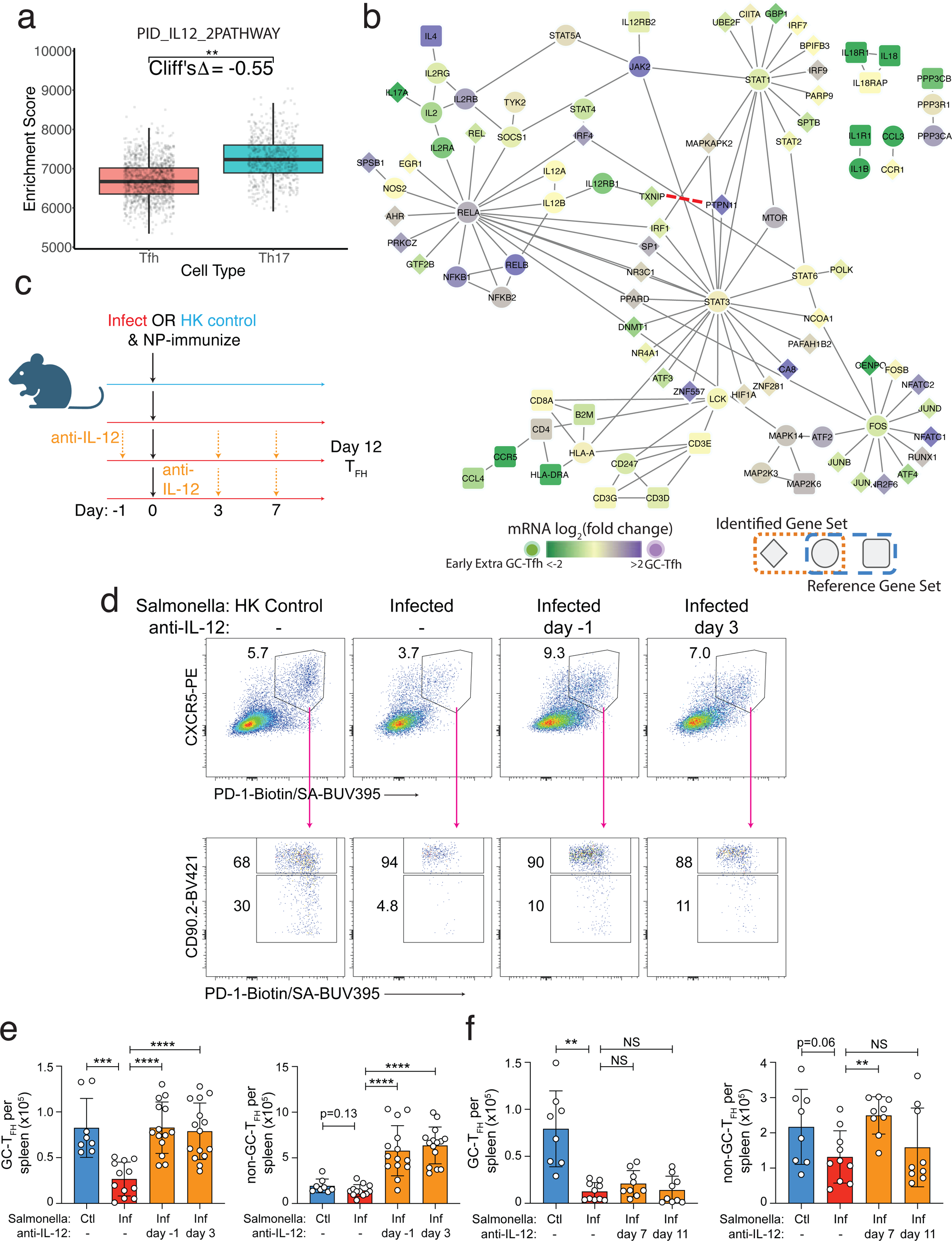
A. scGSEA enrichment of the IL-12 PID pathway pathway for Th17 and Tfh cell populations. Cliffs Δ = 0.56, p-value < 0.0001. B. PID IL-12 pathway protein protein interaction network visualization with the Tfh prioritized gene set overlaid. Red dashed line indicating interaction identified in submodule 1 (Figure 1). Shape and color as previously described. **c-f**, B18+/-mice were given either heat killed (HK) STm control or STm infection, and immunized with NP-CGG in alum on day 0. Mice were additionally treated with neutralizing anti-IL-12 antibodies (or vehicle control) beginning at day -1 or 3 (**e**), or day 7 or 11 (**f**). Tfh were analyzed at 12 days after infection/immunization. C. Experimental design for panels **d** and **e**. D. Representative FACS plots pre-gated on [singlet, live, CD45R-, CD4+, FoxP3-, CD44hi] activated T conventional cells. Shown are gates for total Tfh (CXCR5^high^ PD-1^high^), and GC Tfh (CD90^low^) or non-GC Tfh (CD90^high^). E. Numbers of GC Tfh and non-GC Tfh per spleen. F. Numbers of GC and non-GC Tfh were quantified 12 days after infection as in **d**. For **e-f** charts, circles represent individual mice and bars the mean +/-SD. NS, not significant, * p<0.05, ** p<0.01, ***p<0.001, ****p<0.0001. P values were calculated using two-tailed t tests.

Consistent with prior publications^19^, total Tfh were suppressed; here we additionally and critically show that this effect is primarily driven by the selective suppression of GC Tfh (Extended Figure 5B, C). Indeed, compared to HK STm control, STm infection strongly suppressed the formation of GC Tfh, but minimally affected non-GC Tfh (Figure 6E, Extended Figure 5B). Blocking IL-12 at either day -1 or day 3 fully restored GC Tfh quantity (Figure 6E). Blocking IL-12 also enhanced non-GC Tfh, even though STm infection didn’t suppress non-GC Tfh, potentially suggesting further heterogeneity among non-GC Tfh or additional roles of IL-12.

Given that the effect of blocking IL-12 at day -1 or day 3 was similar, and that both of these times are prior to the formation of bona fide GC T and B cell populations, we next asked whether blocking IL-12 after GCs have developed can still rescue GC Tfh formation. For this, we administered blocking antibodies once at day 7 or 11 after infection, both times that we have previously shown Tfh are already suppressed^19^. Strikingly, blocking at these later times did not rescue GC Tfh formation, indicating that suppression of GC Tfh is not reversible. However, blocking at day 7 did somewhat enhance the non-GC Tfh population (Figure 6F), indicating that there is an early window that extends to day 7 in which IL-12 also inhibits non-GC Tfh development, also consistent with a similar effect in the earlier blocking window experiments (Figure 6E). Collectively, our experimental analysis demonstrates that the effects of IL-12 differ with respect to time and stage of Tfh differentiation, confirming and validating a key and novel aspect of our systems analysis. We also confirm that the IL-12-family member implicated in suppression of Tfh development is IL-12 itself.

## Discussion

Our systems-level approach provides a new perspective on both established and previously unrecognized mechanisms underlying Tfh cell differentiation. A high-quality, well-designed bulk dataset forms the foundation of our analysis, enabling the identification of meaningful biology. We refine these findings using scRNAseq datasets. Critically, we validate key predictions in an in vivo murine model, demonstrating physiological relevance. While traditional differential expression analysis has been the standard for interpreting RNA-seq datasets, this approach often fails to capture complex regulatory relationships. In contrast, network-based approaches incorporate the topology of biological interactions to reveal hidden structure in the data, identifying functionally important genes that may not exhibit strong differential expression on their own. Using a PPI network to incorporate higher-order relationships, additive effects, and functional dependencies among the genes allows us to capture a stronger biological signal. Genes involved in Tfh cell differentiation are more likely to encode interacting proteins that form coherent network submodules. In addition, network propagation serves to filter out genes that may exhibit similar epigenetic signatures but are unlikely to play a functional role in Tfh biology. Critically, this highlights the power of combining several data types through network analysis. Even if the gene encoding the TF is not highly expressed, it can be identified as a critical regulator of the Tfh program.

While we sought to identify key regulators of Tfh differentiation, it is notable that the top-prioritized modules are conserved across species when using this approach. Previous studies have highlighted the significance of several of these modules, but these studies were limited to assessing Tfh in isolation, either using isolated data modalities, in vitro systems, or either human or murine systems. A significant advancement of this work is that the predicted prioritizations are enhanced by combining modalities, not diluted. For instance, the core Tfh modules identified from human tonsil samples also recapitulated known regulators of Tfh differentiation in mice, suggesting the core Tfh gene set is conserved. Our analysis also unifies the previously identified modules by illustrating how these seemingly individual networks are interconnected at the protein-protein interaction (PPI) level.

Fully differentiated GC-Tfh cells from humans and mice share similar surface protein phenotypes and nearly identical gene expression profiles. However, prior in vitro studies attempting to induce Tfh differentiation have often suggested that there are functional differences between species. However, as the methods available for human studies are limited to ex vivo analysis and in vitro approaches, the cells generated in vitro may not correspond to bona fide in vivo GC-Tfh in particular. Thus, they may not fully reflect the differentiation of GC-Tfh populations and so apparent differences between mice and humans may reflect differences in systems and limitations of analysis, rather than true species differences. In vitro systems may not be able to recapitulate complex cognate interactions with GC B cells that GC-Tfh rely on for their differentiation and maintenance. Recent studies demonstrate that DC-mediated interactions can sustain non-GC-Tfh, but not GC-Tfh (PMID). Our network approach, however, highlighted conserved regulators between species, enabling us to formulate hypotheses about Tfh heterogeneity from human data sets and directly test them at a mechanistic level using murine models.

An intriguing candidate emerged from this perspective: the IL-12 signaling pathway. Our analysis of human T cells ranked it as highly important but downregulated among GC-Tfh compared to other pre-Tfh states. Several prior studies utilizing in vitro differentiation of human T cells have reported that IL-12 can enhance or even be necessary for human Tfh differentiation. However, it has also been shown that IL-12 can suppress Tfh and GC formation in multiple murine models of infection. Combining these observations with our own data, we hypothesized that IL-12 may enhance early stages of Tfh differentiation, but block progression to the fully mature GC-Tfh state. To test this, we examined the recently defined murine GC-Tfh and found that IL-12 strongly suppresses their advanced differentiation. Interestingly, non-GC-Tfh were minimally repressed by STm infection, although blocking IL-12 did enhance their differentiation, potentially through other inflammatory signals induced by STm infection. In vitro differentiation, therefore, likely represents an immature Tfh differentiation state similar to pre-Tfh, and for both human and murine cells IL-12 signaling blocks the most mature stage of GC-Tfh differentiation.

Intriguingly, Yeh et al.^33^ demonstrated that GC-Tfh development relies on B cell antigen presentation, while non-GC Tfh can be generated via interaction with DCs. The initial observation that IL-12 could produce Tfh-like cells in vitro in human cells was through co-culture with DCs^17^, so B cell-derived signals could be a missing link for the in vitro differentiation of GC-Tfh. Thus, at present, in vitro systems fail to accurately represent ex vivo GC-Tfh, and studies such as ours, combining high quality human cell analysis with murine experiments, remain crucial for dissecting molecular mechanisms and cellular interactions that are difficult to model in vitro.

In addition to IL-12 signaling, our analysis predicted several modules that have been demonstrated to play roles in Tfh differentiation, but which have been less well-studied. One intriguing protein is PTPN11 (SHP-2). PTPN11 has been implicated in PD-1 signaling in GC-Tfh, and PD-1 contributes to the localization of Tfh within the GC. Consistent with this and human data^57^, GC-Tfh were the highest expressors of PD-1 (Fig. 6d), and our network connects PTPN11 and PD-1 (PDCD1, Fig. 2d). However, the molecular mechanism of SHP-2 function in this was not investigated. Other modules, such as HIF1A, have mixed reports of their role in Tfh differentiation^73–75^. Our analysis predicts that Hif1a is important but is not differentially expressed between non-GC-Tfh and GC-Tfh (Fig. 2g). IRF4 was identified (Fig. 2g), which has been shown direct Tfh versus Th1 differentiation through reading out TCR signaling strength^76^. Consistent with this, our analysis shows IRF4 connected to NFATC2 and a hub of other molecules that regulate TCR signaling (Fig. 4e). While some reports highlight the importance of TCR signaling in Tfh^57,76–80^, our analysis reveals a sub-network around the NFATC2 hub, suggesting that intriguing and important biology remains to be discovered in how TCR signaling regulates Tfh differentiation.

The integrative analysis reported here succeeds in capturing and encompassing over a decade of Tfh-related discoveries while at the same time revealing multiple novel candidate genes and pathways. Although curated gene sets may help define Tfh identity in some contexts, they often lack the ability to resolve cross-context differences. The prioritized gene sets identified in this work converge on a core regulatory network that is conserved across species, tissues, disease states, and experimental systems, an outcome only possible if the underlying biology is truly consistent. The identified modules and genes offer a rich resource for future functional studies and therapeutic development. Notably, our findings help resolve a long-standing debate in the field regarding the necessity of IL-12 for Tfh differentiation, with important implications for vaccine strategies that utilize IL-12–based adjuvants. Further, this method could be applied to other biological contexts outside of Tfh differentiation and provides a roadmap for dissecting immune cell differentiation which has broad relevance for immunotherapy and vaccine design.

## Figure Legends

**Extended Figure 1.**
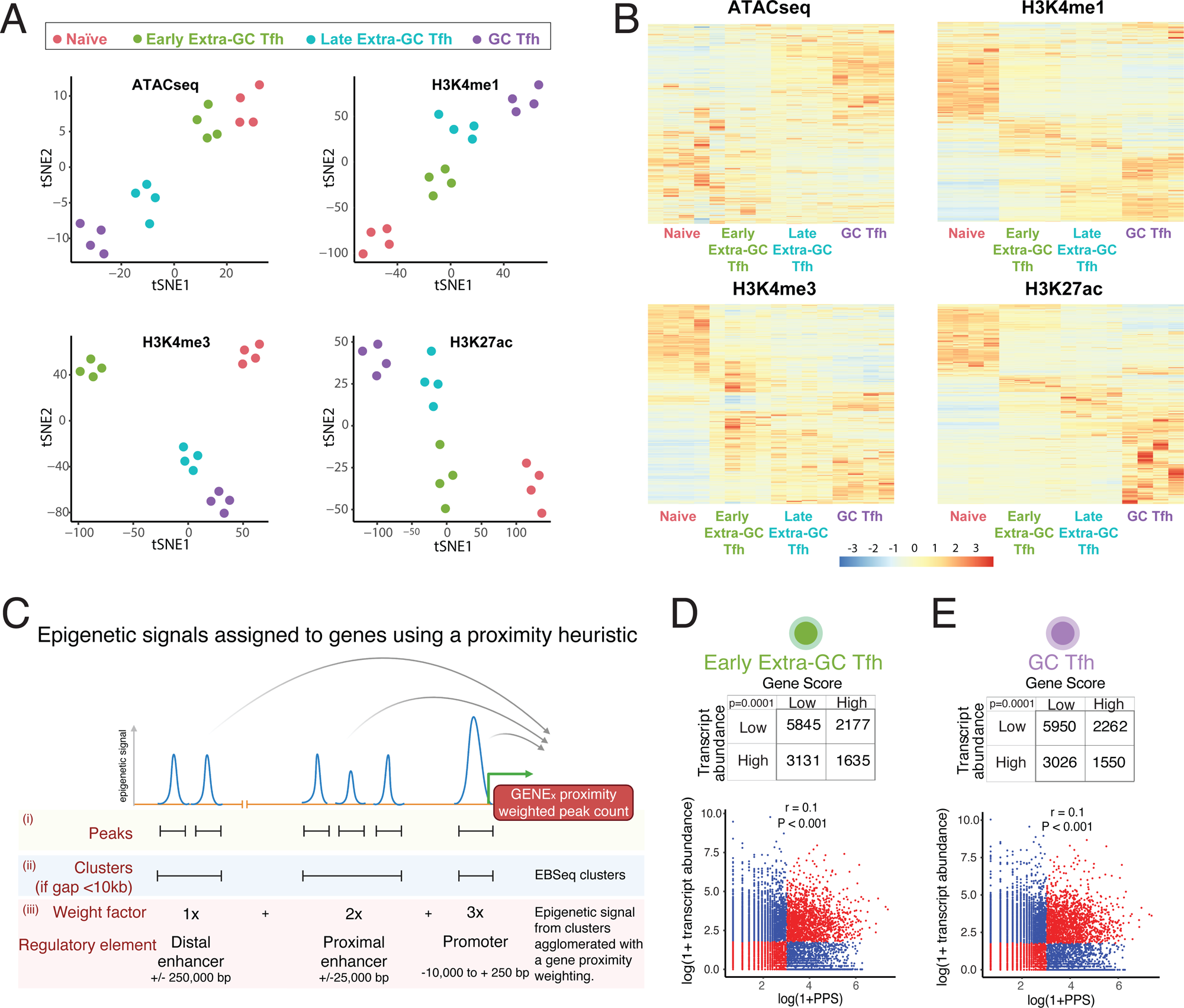
A. t-SNE visualizing epigenomic profiles (ATAC-seq, H3K4me1, H3K4me4, H3K27Ac) of samples at different stages in the Tfh cell differentiation trajectory. B. Heatmap visualizing differentially regulated regions for each of the 4 epigenomic datasets - ATAC-seq, H3K4me1, H3K4me4, H3K27Ac C. Schematic of a proximity-weighted count heuristic to calculate peak proximity scores. D. Relationship between PPS and transcript abundances at the Early Extra-GC Tfh state in Tfh differentiation (PPS threshold 20, transcript abundance threshold 5) E. Relationship between PPS and transcript abundances at the GC Tfh state in Tfh differentiation (PPS threshold 20, transcript abundance threshold 5)

**Extended Figure 2.**
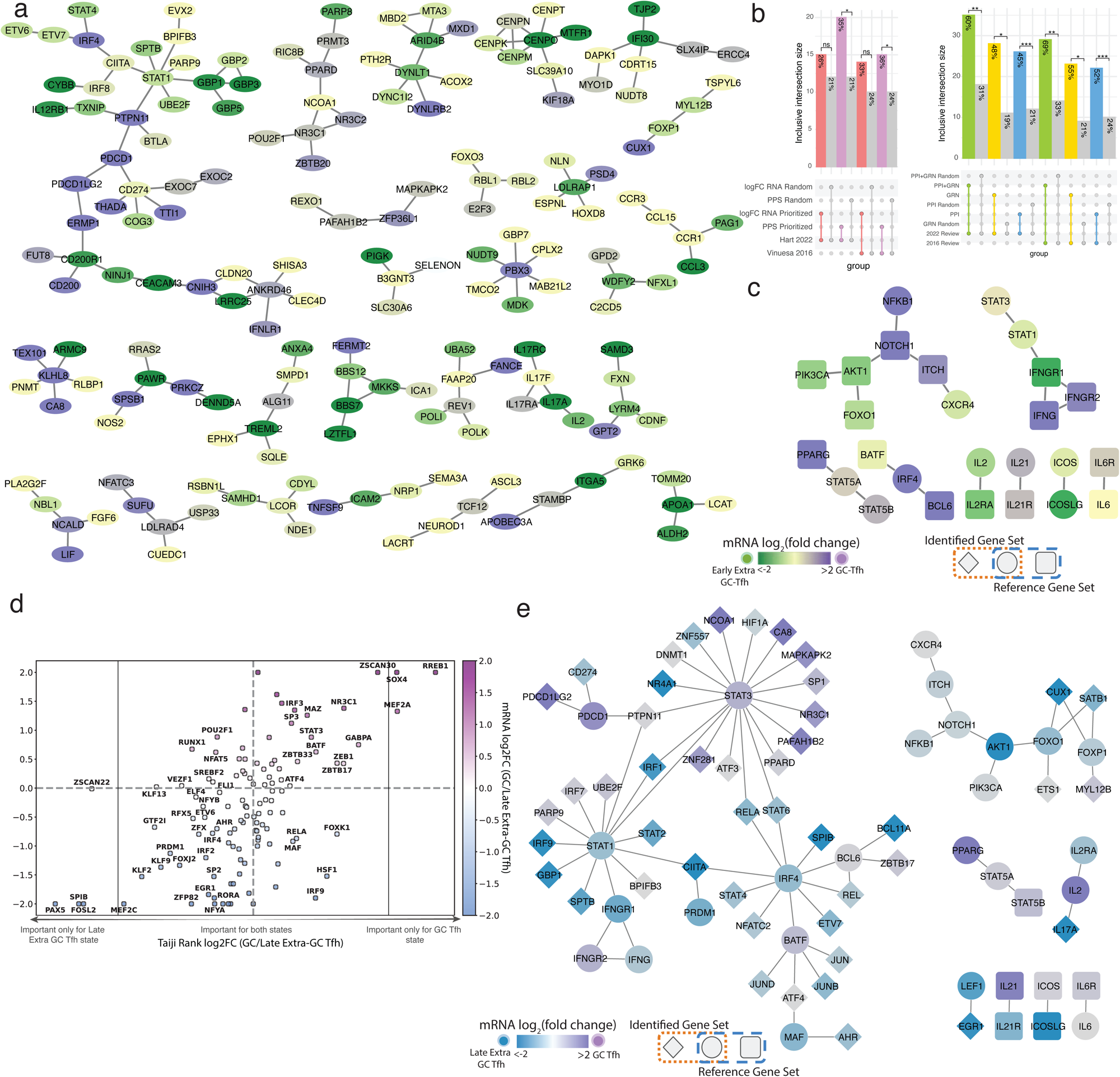
A. All 27 modules identified with the protein-protein interaction network propagation colored by the mRNA log_2_FC of the GC and Early Extra-GC Tfh where higher expression in GC is shown in purple, and higher in Early Extra-GC Tfh is shown in Green. B. Overlay of the logFC RNA prioritized set with direct interactors on the Vinuesa 2016 reference network. Shared genes (circle) and reference only genes (square) are covered by mRNA log_2_FC as previously described. C. Overlap of size matched gene sets of log_2_FC RNA (left) and top PPS (right) with their first-degree interactors with 2 reference Tfh sets. Size matched sets (184 genes) of the top log_2_FC RNA scores or PPS were identified and first degree interactors were added from the network. 1000 permutations of a random size matched sets (184 genes) with first degree interactors added were also compared for each analysis. Bi-nomial proportions test performed with average random overlap and average random set size, P-value threshold <0.05. D. A focused scatter plot on the driver TF set without ZNFs analyzed by Taiji given by the mRNA log_2_FC between the GC and Late Extra GC state (y-axis) and the Taiji Rank log_2_FC between the 2 states (x-axis) and colored by the mRNA log_2_FC of the GC and Late Extra-GC Tfh where higher expression in GC is shown in purple, and higher in Late Extra-GC Tfh is shown in Blue. E. Overlay of the PPI+GRN prioritized set with direct interactors on the Vinuesa 2016 reference network colored by mRNA log_2_FC of the GC and Late Extra-GC Tfh where higher expression in GC is shown in purple, and higher in Late Extra-GC Tfh is shown in Blue. Shared genes (circle), analysis only discovered interactors (diamond), and reference only genes (square).

**Extended Figure 3.**
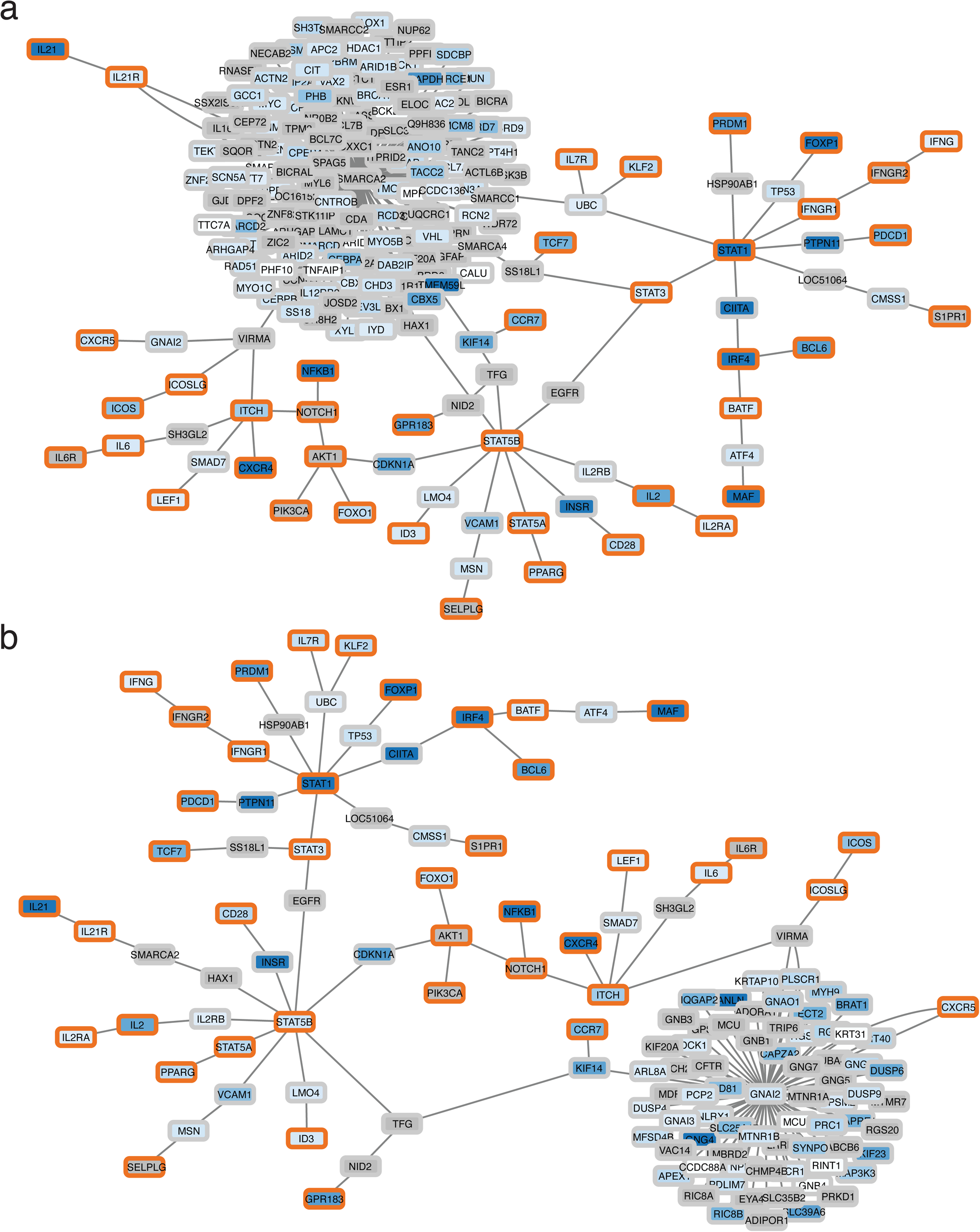
A. IL21 connection to the rest of the Vinuesa genes through the SMARCA2 hub in the PPI network. The shortest path connecting all Vinuesa genes was used with all interactors of SMARCA2 visualized. Orange, in Vinuesa genes; grey, gene in the PPI network. B. CXCR5 connection to the rest of the Vinuesa genes through the GNAI2 hub in the PPI network. The shortest path connecting all Vinuesa genes was used with all interactors of GNAI2 visualized. Orange, in Vinuesa genes; grey, gene in the PPI network.

**Extended Figure 4.**
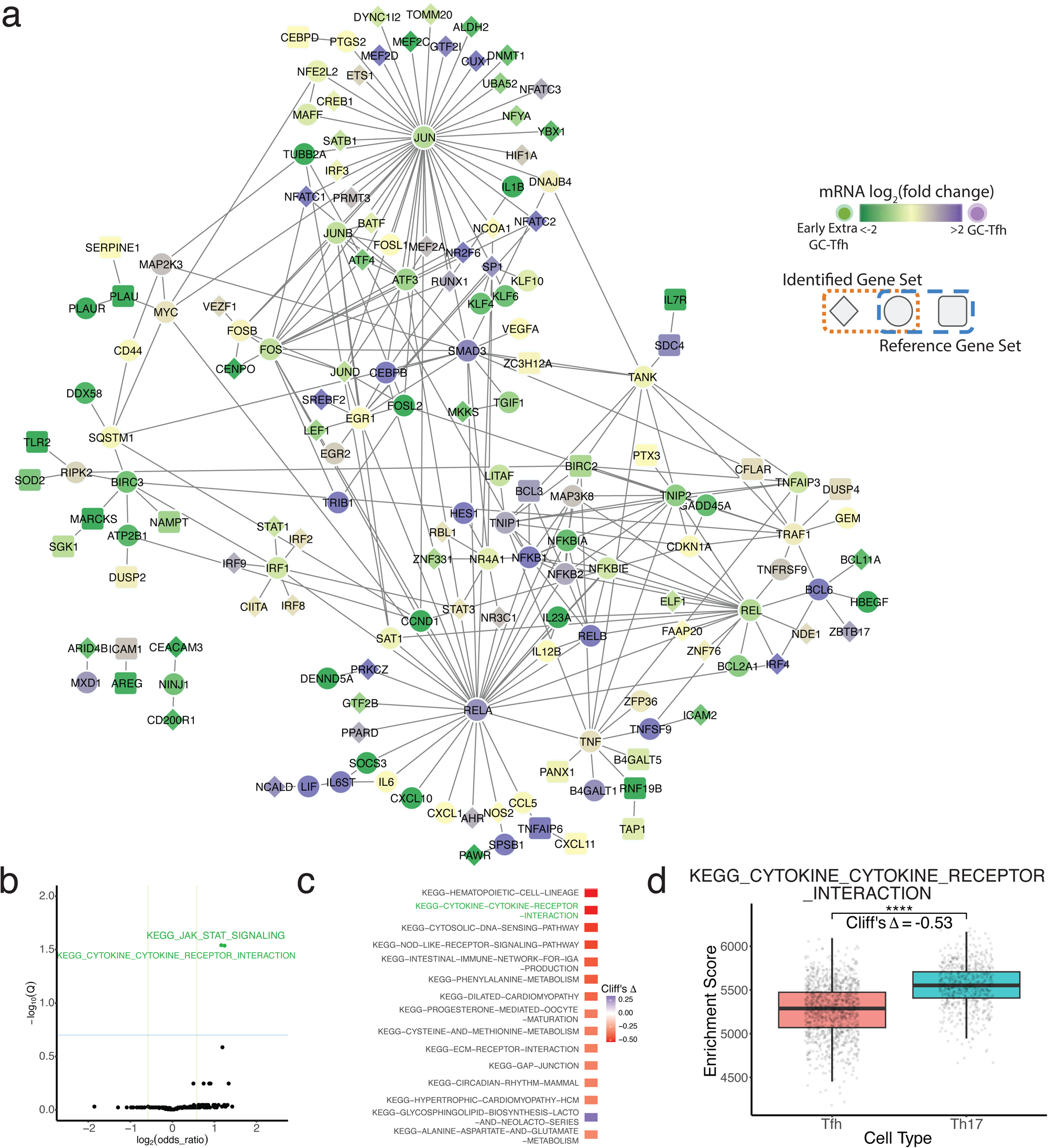
A. HALLMARK TNFA-SIGNALING-VIA-NFKB pathway PPI network visualization with the prioritized Tfh gene set overlaid. Shape and color as previously described. Shared genes (circle), analysis only discovered interactors (diamond), and reference only genes (square) are colored by mRNA log_2_FC as previously described. B. Volcano plot visualizing effect sizes and significance of functional overlaps between the Tfh identified gene set and KEGG pathways. Significant pathways colored in blue. Significant pathways colored in blue (-log(Q-value) > 0.7, log2(odds_ratio) > 0.58). Adjusted p-value (FDR) < 0.2, and 1.5-fold enrichment. C. Human scRNA top 15 KEGG pathways enriched for the Tfh identified gene set given by the top effect size between the Tfh and Th17 cell population enrichment scores. D. scGSEA enrichment of the CYTOKINE-CYTOKINE-RECEPTOR-INTERACTION KEGG pathway for Th17 and Tfh cell populations (Cliffs Δ = -0.55, p-value < 0.01).

**Extended Figure 5.**
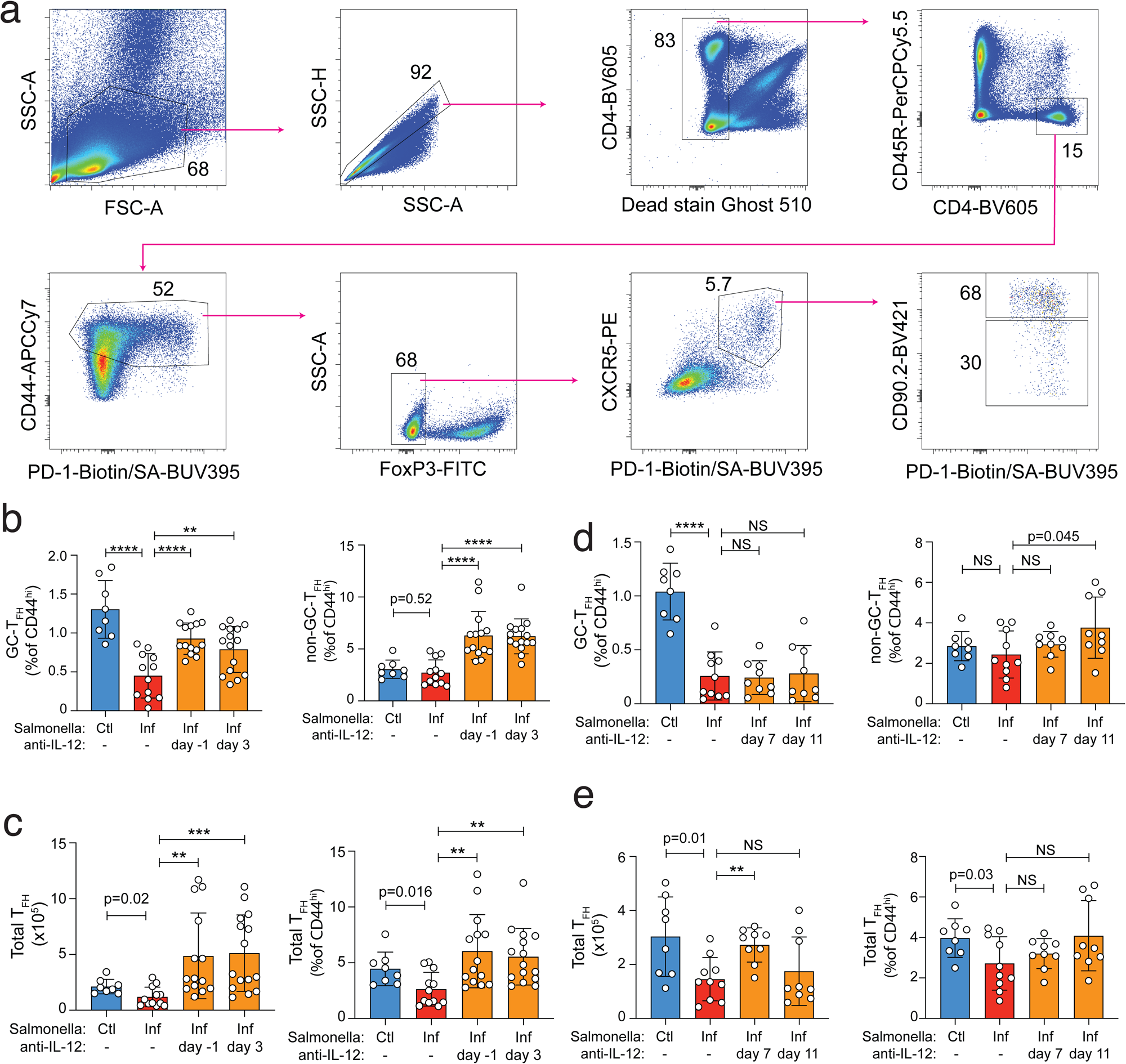
**a-c**, B18+/-mice were given either heat killed (HK) STm control or STm infection, and immunized with NP-CGG in alum on day 0. Mice were additionally treated with neutralizing anti-IL-12 antibody beginning at day -1 or 3 (**a-c**), or day 7 or 11 (**d,e**). Tfh were analyzed at 12 days after infection/immunization. A. Representative FACS plots illustrating gating used to identify GC and non-GC Tfh among activated T conventional cells. B. Percentage of GC (CD90^low^) or non-GC (CD90^high^) Tfh (CXCR5^high^ PD-1^high^) among activated T conventional cells, following IL-12 block at day -1 and 3. C. Number (left) and percentage (right) of total Tfh (CXCR5^high^ PD-1^high^) following IL-12 block at day -1 and 3. D. Percentage of GC (CD90^low^) or non-GC (CD90^high^) Tfh (CXCR5^high^ PD-1^high^) among activated T conventional cells following IL-12 block at day 7 or 11. E. Number (left) and percentage (right) of total Tfh (CXCR5^high^ PD-1^high^) following IL-12 block at day 7 or 11.

## Methods

### Human Flow cytometry and sorting

600 million cells from healthy adult human tonsils were resuspended in Miltenyi magnetic-activated cell sorting (MACS) buffer and stained with biotinylated anti-human CD19 (clone: HIB19; BioLegend, #555411) on ice for 30 min. Cells were washed once in MACS buffer and incubated with antibiotin microbeads (Miltenyi Biotec) for 30 minutes on ice. Cells were then loaded on Miltenyi LS columns at 300 million cells per column, and the flow-through was collected as the B cell–depleted fraction. These cells were spun down and subjected to Fc block (Human BD Fc Block BD Biosciences, #564219), washed with 1% BSA in PBS, and stained with LIVE/DEAD Fixable Blue Dead Cell Stain (ThermoFisher Scientific, #L23105), before they were surface stained with antibodies against CD3 (clone: SK7; BioLegend, #344817), CD4 (clone: OKT4; BioLegend, #317420), CD45RA (clone: H100; BioLegend, #304122), CXCR5 (clone: J252D4; BioLegend, #356920), and PD1 (clone: EH12.2H7; BioLegend, #329924). The following populations were sorted using FACSAria2 (Becton Dickinson), as shown in the gating strategy present in Figure 1C: naive CD4 T cells (CD4+CD45RA+CXCR5-PD1-), (ii) Early Extra-GC Tfh (CD4+CD45RA-CXCR5+PD1-), (iii) Late Extra-GC Tfh (CD4+CD45RA-CXCR5+PD1+), and (iv) GC Tfh cells (CD4+CD45RA-CXCR5++PD1++). All antibodies were used at a dilution of 1:40.

### Bulk RNA-seq and analysis

Total RNA was isolated from the FACS-sorted healthy tonsillar cells using the RNeasy plus Micro Kit, according to the manufacturer’s instructions (Qiagen). RNA-sequencing libraries were prepared as previously described ^81^. Briefly, whole transcriptome amplification and tagmentation-based library preparation were performed using the SMART-Seq2 protocol. Libraries were assessed for quality using High Sensitivity DNA chips on the Agilent Bioanalyzer, quantified using Qubit fluorometer (Thermo Fisher Scientific), as well as KAPA Library Quantification kit (KAPA Biosystems), followed by 35-bp paired-end sequencing on a NextSeq 500 instrument (Illumina). 5–10 million reads were obtained from each sample and aligned to the University of California, Santa Cruz hg38 transcriptome. Gene expression was calculated using RSEM as previously described (Li & Dewey, 2011), with alignments to ENSEMBL hg38, and differential expression analysis was conducted using EBSeq, with differentially-expressed genes determined within each pattern by selecting genes with a PPDE (posterior probability of differential expression) of at least 0.95.

### CUT&RUN

CUT&RUN libraries were prepared using rabbit monoclonal antibodies against H3K4me1, H3K4me3 and H3K27Ac (Cell Signaling Technology catalog numbers 5326, 9751, and 8173, respectively) as previously described. The protein-A Mnase fusion protein was a gift from Steven Henikoff.

### ATAC-seq

ATAC-Seq libraries were prepared as originally described^35^. Fifty thousand freshly sorted cells of each population were pelleted and washed with 50 μL chilled 1× PBS, and with 50 μL lysis buffer (10 mM Tris-HCl pH 7.4, 10 mM NaCl, 3 mM MgCl2, 0.1% IGEPAL CA-630). The nuclei were pelleted in a cold micro-centrifuge at 550×g for 10 min, and resuspended in a 50μL transposition reaction with 2.5μL Tn5 transposase (FC-121-1030; Illumina) to tagment open chromatin. The reaction was incubated at 37°C in a Thermomixer (Eppendorf) at 300 rpm for 30 min. Tagmented DNA was purified using a QIAGEN MinElute kit and amplified with 7 or 11 cycles of PCR, based on the results of a test qPCR. ATAC-Seq libraries were then purified using a QIAGEN PCR cleanup kit and quantified using KAPA library quantification kit (KAPA Biosystems, Roche) and sequenced on the Nextseq 550 platform.

### Human scRNA-seq datasets

Single cell RNA sequencing (Chromium Next GEM Single Cell Gene Expression, 10X Genomics) was done. Thawed cells were counted, and the viability was determined by staining the cells with Trypan blue. Isolated CD45+ cells were centrifuged and resuspended at a concentration of 800– 1200 cells per microliter in RPMI with 2% (v/v) human AB serum and loaded onto 10X Chromium instrument and Single Cell Gene Expression (GEX) libraries were prepared with the following kit: Chromium Single Cell 5’ V1 (10X Genomics). 10,000 cells were targeted for each sample. Isolated cells were transposed and partitioned into Gel Beads-in-emulsion (GEMs) using the 10x Chromium Controller and Next GEM Chip J. GEMs were visually inspected, and only samples with an opaque and uniform aspect were used for library preparation. Representative traces and quantitation of libraries were determined using Bioanalyzer High Sensitivity DNA Analysis (Agilent, Santa Clara, CA). Sequencing was done on Illumina NovaSeq S4 targeting 20,000 reads per nucleus for gene expression.

Fastq files were processed 10x Genomics Cell Ranger v5.0.1 using 10x Genomics Cloud Analysis^82,83^. Reads were mapped to the GRCh38 human reference genome and counted without depth normalization. The filtered count matrix was then analyzed using the Seurat (v5.1.0)^84^ in R. Low quality cells were identified and removed based on the following criteria: more than 100 RNA unique molecular identifiers, less than 25% mitochondrial read fraction. Doublet cells were identified using the DoubletFinder package in R^85^. Only CD4+ population cells were analyzed further.

We performed single-cell RNA-seq analysis using Seurat ^86^ (v5.0.0) R version (4.4.0), followed by batch correction with Harmony to integrate data across experimental conditions. Outlier cells were removed prior to analysis to ensure high-quality clustering. Genes associated with mitochondrial content, ribosomal components, hemoglobin, sex chromosomes, and the surfactant protein family were excluded to reduce technical and tissue-specific bias. Gene expression values were then normalized. We identified the top 5,000 highly variable features using the variance-stabilizing transformation (vst) method, and scaled gene expression values across cells. Principal component analysis (PCA) was performed using these variable genes, and the resulting principal components were corrected for batch effects using the Harmony algorithm. An elbow plot of the Harmony components was used to determine the number of informative dimensions, and the first 20 components were selected for downstream analysis. Clustering was performed using shared nearest neighbor (SNN) graph construction and the Louvain algorithm, with parameters k.param= 50 and resolution = 0.7 based on the Harmony-reduced space.

We identified cluster 2 as primarily Tfh cells by using a composite expression score of Tfh markers: BCL6, CXCR5, CXCL13, CD200, PDCD1. Further, the Azimuth Human Tonsil v2 reference^87^ was used to label the gut CD4+ cells. The cells in the overlap of cluster 2 with either Azimuth GC-Tfh-SAP or Tfh T:B border labels were chosen for the Tfh group.

We also identified 2 non-Tfh populations for comparison: Th17 and Central Memory (CM) CD4+. First, we removed all Tfh, Naive, Cycling, Treg/Eff-Tregs-IL32, and Tfh adjacent (Tfh mem, GC-Tfh-OX-40 and Tfh-LZ-GC) labeled cells from the dataset. A composite score of Th17 markers (CCR6, IL23R, and RORC) was used to select cluster 5 as the Th17 cells. For the CM non-Tfh CD4+ group, the dataset was reclustered using the Azimuth Human PBMC reference ^88^ and used markers to visualize the cells (CCR7, SELL). The cells labeled central memory and effector memory were selected. For all groups, only cells from people living with HIV (PLWHIV) were kept for analysis.

### Murine scRNA-seq datasets

We processed publicly available mouse lymph node scRNA-seq data from Kun et al.^95^ allergy model. We followed a standard Scanpy^96^ (v1.10.3) workflow. Raw counts were normalized per cell and log-transformed with default parameters. Top 2000 highly variable genes were selected. We performed principal component analysis (PCA) and embedded neighborhoods in UMAP space. Potential cell doublets were identified and removed using Scrublet^97^ implemented in Scanpy with an expected doublet rate of 0.05.

Major immune cell types were annotated based on the expression of canonical marker genes: Cd19 and Ms4a1 for B cells; Cd3d and Cd4 for CD4+ T cells; Cd3d and Cd8a for CD8+ T cells; Klrb1c and Klrk1 for NK cells; Itgax for DCs. Cells co-expressing *Ms4a1* (CD20, B cell marker) and *Cd3d* (T cell marker) were identified as potential T/B doublets and removed. We further subsetted the CD4+ T cell compartment and identified CD4+ T cell subtypes using established lineage-defining markers: Bcl6 and Pdcd1 for Tfh; Gata3 and Ccr4 for Th2; Il2ra and Foxp3 for Treg. CD4+ T cells negative for Tfh, Th2, and Treg markers and exhibiting low expression of activation markers (Cd69, Icos) were designated as resting CD4+ T cells. scGSEA was performed on the Tfh and Resting CD4+ T cell populations using the escape R package to identify the cell enrichment score for each gene set of interest and cell type.

### Human spatial datasets

Cells with Tfh or CD4+ T-cell labels were extracted from human tonsil slide-tag snRNA-seq data ^59^ with the R package Seurat. From the CD4+ T-cell group, naive T-cells were filtered out using a composite score of the following markers (‘CD3E’, ‘CCR7’, ‘LEF1’, ‘SELL’, ‘GATA3’, ‘CD4’, ‘LTB’, ‘RCAN3’, ‘CD40LG’, ‘LTB’, ‘PIK3IP1’, ‘TCF7’) with a threshold of >10. scGSEA was performed using the escape R package to identify the cell enrichment score for each gene set of interest and cell type. The cells were visualized in 2D space and colored by enrichment.

### Personalized PageRank identification of driver TFs

To identify candidate driver transcription factors (TFs), we applied the personalized PageRank algorithm implemented in Taiji v1.2.0^41^. A sample configuration file is provided. Briefly, both epigenomic datasets (ATAC-seq, H3K4me1, H3K4me3, H3K27Ac) and bulk RNA-seq data were organized by biological replicate and by cell state (Naive, Early Extra-GC Tfh, Late Extra-GC Tfh, and GC Tfh).

A separate Taiji run was performed for each of the four epigenomic modalities, each paired with the bulk RNA-seq dataset. For the ATAC-seq runs, input consisted of four replicate pairs of narrowPeak and BED files per state, with the file format specified as ‘NarrowPeak’. For the CUT&RUN datasets (H3K4me1, H3K4me3, H3K27Ac), four replicate BAM files were used per state, with the tag set to ‘PairedEnd’. In all runs, four replicate TSV files of bulk RNA-seq per state were provided, with the input tag set to ’GeneQuant’. The human genome assembly GRCh38 was used throughout.

Taiji’s output GeneRanks files were filtered to retain only genes with a p-value < 0.01. Within each epigenomic dataset, genes were ranked per state, and for each gene, the minimum rank across the four states was determined. The 100 genes with the lowest minimum ranks were selected from each dataset. These top-ranked TFs were then combined across the four datasets, yielding a final set of 152 unique transcription factors. To visualize the results, the Taiji rank was averaged across the 4 datasets and the log2FC between the average GC and Early Extra-GC Tfh or Late Extra-GC Tfh Ranks or Expression were calculated.

### Calculation of gene-centric peak proximity scores

For the chosen set, peak proximity scores (PPS’s) for a gene (“genecentric PPS”) were calculated by computing a weighted sum of the number of peak clusters from the pattern in the gene’s vicinity using distance-based weights to reflect the varying contributions of promoter (± 10 kb) [3x weight], proximal enhancer (± 25 kb) [2x weight] and distal enhancers (± 250 kb) [1x weight] to gene regulation.

Gene-centric PPS scores were first calculated individually for each epigenomic dataset. These were then combined into a final PPS score using equal weights for the different datasets.

### Network propagation using random walk with restart

To integrate PPS scores, we used network propagation based on the random walk with restart algorithm, implemented in HotNet2^54^. The HotNet2 code is available at https://github.com/raphael-group/hotnet2. Propagation was performed on the union of high-quality reference human binary and co-complex protein-protein interaction networks from the HINT database^53^.

The input influence matrix of the input network, permuted networks, and their corresponding influence matrices which are required for network propagation were prepared using the makeNetworkFiles.py script provided with HotNet2. For a robust analysis, we generated 500 permuted networks and 500 heat vector permutations. All files were saved in HDF5 format.

Network propagation was run using the default HotNet2 parameters: beta = 0.4 and four delta values automatically selected by the software. A sample configuration file is provided. Modules were considered significant if they had a P-value < 0.05 and contained more than three genes. We performed five independent runs and retained only the subnetworks that appeared consistently in all five runs. These reproducible subnetworks are referred to as the PPI prioritized gene set.

The PPI prioritized networks were visualized using Cytoscape ^90^ v. 3.10.2 and colored according to log fold change (logFC) values across the differentiation trajectory, where fold change was calculated as log2(GC Tfh / Early Extra-GC Tfh). A random sample of 2,000 edges from the reference PPI network was also visualized alongside the subnetworks using the R package qgraph^91^.

The NetworkX (v. 2.5) ^92^ steiner_tree python package to identify the shortest path between the genes in the 2016 reference set from the PPI network. Then all interactors of a gene of interest were added to the graph and visualized in Cytoscape. Genes belonging to the 2016 reference set were outlined in orange and the node color corresponds to the node PPS score prior to the network propagation with dark blue being the highest expression and grey indicating no value.

### Functional overlap between Tfh cell subnetworks and published datasets

We evaluated the functional overlap between our identified Tfh gene sets and published Tfh gene sets, where functional overlap was defined as either exact gene matches or genes that directly interact. Established Tfh gene sets were derived from three reviews: Vinuesa et al. (2016) ^55^, Hart and Laufer (2022)^56^, and a review of mouse Tfh genes by Crotty et al (2020)^23^. We compared the functional overlap of genes prioritized by our combined network propagation approach to those identified by protein interaction network propagation alone, Taiji alone, and to the average overlap from 1,000 random network-derived gene sets matched in size for each dataset. To test the importance of network propagation, we also selected top-scoring genes based solely on individual gene-level metrics. A size-matched set of the PPI prioritized gene set (184 genes) top log_2_FC between GC Tfh and Early Extra-GC Tfh was used for RNA-seq data and PPS scores was used for epigenomic data. Functional overlap for these gene sets was similarly compared to 1,000 total gene set size-matched random gene sets and average random overlap was calculated. P-values were calculated using a Fisher’s exact test. A second version of creating the random baseline was also tested. 1000 size-matched sets to the prioritized genes (184) were randomly generated and their direct interactors were included. This led to a varying size of the random set so the average random overlap and average random gene size were used for the binomial proportions test p-value calculation.

The PPI network overlap between the reference Tfh gene sets and the identified Tfh gene set was created by extracting the corresponding nodes and edges from the HINT database associated with the reference genes, as well as including additional direct interactors from the prioritized Tfh gene set. The PPI network overlap was visualized using Cytoscape. We colored these overlapping modules based on the logFCs across the trajectories, where fold change was defined as log_2_(GC Tfh/Early Extra-GC Tfh) or log_2_(GC Tfh/Late Extra-GC Tfh).

### scRNA enrichment of Tfh cell subnetworks and published datasets

For the scRNAseq analyses, scGSEA was performed using the escape R package ^93^ to identify the cell enrichment score for each gene set of interest and cell type. Four gene sets were evaluated: two references derived from published review articles, one composed of Tfh cell subnetwork genes, and one containing the driver TFs selected by Taiji. Enrichment scores were compared across cell types to assess differential pathway activity. Statistical significance was determined using p-values calculated from the Wilcoxon rank-sum test and effect sizes were quantified using Cliff’s Δ to capture the magnitude of enrichment differences. The data was visualized using boxplots with a dot overlay. Only 70% of the population was represented by dots and was randomly sampled. Functionally different datasets between cell types were considered significant by passing both Wilcoxon rank-sum test p-value (< 0.05) and effect size given by the Cliffs Δ ≥ abs(0.1).

### Pathway enrichment calculations

Pathways were extracted using the msigdb R package (v7.5.1)^94^. Top pathways were identified for bulk RNA and scRNA datasets from 3 different databases-KEGG, Hallmark, and PID across species and tissues. The PPI network overlap between the pathway gene sets and the identified Tfh gene set was created and visualized as previously described.

For the bulk RNA-seq pathways enriched with the Tfh prioritized gene set were detected by calculating the functional overlap between the Tfh prioritized gene set and all pathways in the 3 different databases. The significance of gene set overlap was assessed by comparing to the average overlap of 1,000 random, size-matched sets of network genes. Functionally overlapping pathways were considered significant if the odds ratio (defined as the odds of overlap between the identified gene set and a given pathway divided by the odds of overlap from a random gene set) exceeded 1.5, and the false discovery rate (FDR) was below 0.2. P-values were calculated using a binomial proportions test and adjusted for multiple comparisons using the Benjamini-Hochberg correction. Significant modules were visualized as described previously, with color representing the log2FC across states.

For the scRNAseq analysis, scGSEA was performed using the escape R package to identify the cell enrichment score for each pathway. Functionally different pathways between cell types were considered significant by passing both Wilcoxon rank-sum test p-value (< 0.05) and effect size given by the Cliffs Δ ≥ abs(0.1). The top 15 pathways with the highest abs(Cliffs Δ) value were reported.

### Murine Salmonella Experiments

#### Mice

B1-8+/-^98^ BALB/c mice, containing elevated frequencies of NP-specific B cells, were used for all mouse experiments. Mice were bred and housed in specific pathogen free conditions in a room at 20-26°C with 30-70% relative humidity. All experiments were conducted under protocols approved by the University of Pittsburgh Institutional Animal Care and Use Committee (IACUC). All mice were between the age of 8-16 weeks at the start of experiments. Males and females were both used and no differences were noted.

#### Bacterial strains

*Salmonella enterica* serovar Typhimurium (STm) *aro*A attenuated strain SL3261^99^, provided by Roy Curtiss III, Arizona State University) was used for all experiments. Glycerol stocks were prepared by growing bacteria overnight in Luria broth, then mixed 1:1 with sterile 50% glycerol and stored at -80°C.

#### Infection, Immunization, and anti-IL-12 Treatment

To prepare STm, one day prior to injection, stabs of frozen STm glycerol stock were inoculated into Luria broth (Fisher) containing 100 mg/mL streptomycin and grown at 37°C overnight with shaking. The day of injection, heat killed bacteria were prepared as follows: cells were pelleted at 1800*g*, washed twice in PBS, resuspended in PBS, transferred to a fresh tube, and incubated at 56°C for 1 hour. For infectious inoculum, bacterial cultures were split 1:25 into fresh media and grown for an additional 2 hours at 37°C to allow bacteria to reach mid-log phase, then washed 3 times in 20°C sterile PBS. Bacterial concentration was estimated by absorbance at OD600, and approximately 5x10^5^ colony forming units were injected i.p. in 200 mL PBS. Colony forming units of infectious inoculum, and absence of live bacteria in heat killed inoculum, were confirmed by overnight growth at 37°C of serial dilutions of inoculum on Luria broth agar (BD Difco) plates with streptomycin. For NP-CGG immunization, mice were immunized with 50 mg NP-CGG precipitated in alum i.p. in 200 mL PBS. The ratios of NP to CGG ranged from 31-33.

For IL-12 blocking experiments, mice were infected with STm or given HK STm, then immunized with NP-CGG 3-5 hours later. All mice were given an initial loading dose of 200 mg anti-IL-12 in PBS by intraperitoneal injection (BioXcell catalog number BE0051, clone C17.8). Mice in day -1 treatment groups received an additional 100 mg at days 3 and 7. Mice in day 3 treatment groups received another 100 mg at day 7. Mice in day 7 and 11 treatment groups only received the loading dose.

#### Flow cytometry analysis for mouse experiments

Spleen pieces were weighed and made into single cell suspensions by mechanical disruption in STm media (PBS + 5% bovine serum + 2 mM EDTA). RBC were lysed with ACK lysis buffer for 60 seconds, washed and resuspended in STm media, and live cells enumerated with trypan blue on a hemacytometer. Cells were pelleted and washed once in PBS before staining. Five million cells per sample were stained at 100 million cells/mL with the dead cell discriminator Ghost Violet 510 (Tonbo, per manufacturers protocol) in PBS, and washed with staining media (SM; PBS with 3% bovine serum (Gibco), 1 mM EDTA, and 0.02% sodium azide). Cells were stained with fluorescently conjugated antibodies as labeled in figures in SM for 20 min on ice, then washed twice with staining media, and fixed with 1% PFA in PBS for 20 minutes on ice. For intracellular stains, cells were instead fixed/permeabilized with FoxP3 Transcription Factor staining Kit (Invitrogen) for 30 minutes on ice. After fixation, all wash steps pelleted cells at 750*g*. Cells were washed twice with FoxP3 Kit permeabilization buffer, resuspended with 25 uL SM containing 10% mouse and 10% rat serum, and 25 mL SM containing 2x concentration of intracellular stain antibodies (CXCR5, T-bet, Bcl-6, and FoxP3) was added, and the mixture incubated overnight at 4°C. Cells were then washed twice with FoxP3 Kit permeabilization buffer, resuspended in SM, and data collected using BD Biosciences LSRII or Fortessa or Cytek Aurora. All wash steps used 500*g* to pellet cells here and for all remaining sections unless otherwise specified.

Antibodies used: CD4-BV605 clone GK1.5, BD Biosciences catalog number 563151, used at dilution 1:400. CD45R-PerCPCy5.5 clone RA3-6B2, Biolegend catalog number 103236, used at dilution 1:50. CD44-APCCy7 clone IM7, Biolegend catalog number 103028, used at dilution 1:200. PD-1-Biotin clone G4, grown and conjugated in house, used at dilution 1:400. SA (streptavidin)-BUV395, BD Biosciences catalog number 564167, used at dilution 1:200. FoxP3-FITC clone FJK-16s, Invitrogen/ThermoFisher catalog number 11-5773-82, used at dilution 1:400. CXCR5-PE clone 2G8, BD Pharmingen catalog number 551959, used at dilution 1:100. CD90.2-BV421 clone 53-2.1, Biolegend catalog number 140327, used at dilution 1:500.

#### Quantification and statistical analysis for murine studies

No statistical methods were used to pre-determine sample sizes, but our sample sizes are similar to those reported in our previous publications. Data collection and analysis were not performed blind to the conditions of the experiments. Mice were assigned to experimental groups based on genotype. The experiments were not randomized. For Salmonella infected mice, mice that showed poor infection as determined by >50x lower than average splenic bacterial CFU at necropsy compared to others in the group were removed. Data distribution was assumed to be normal, but this was not formally tested.

Statistical significance was quantified with GraphPad Prism software (version 10) by two-tailed *t*-test or as indicated in each figure. A P-value <= 0.05 was considered significant, with the following symbols used to denote P-value ranges: p < =0.05(*), p < 0.01 (**), p < 0.001 (***), p < 0.0001 (****). Cell number was calculated per spleen using a combination of FACS analysis and trypan blue live cell counts as follows: the percent of cell of interest among live cells was determined by FACS, then the ratio of spleen processed for FACS analysis by weight as a portion of the total weight was used to calculate the total number of cells of interest per spleen. All flow cytometry data was analyzed using FlowJo version 10 (BD), spectral flow cytometry data was unmixed using SpectroFlo (version 3, Cytek Biosciences), and figures prepared using Adobe Illustrator version 2025.

## Code and Data Availability

All datasets and documentation are available via GitHub at https://github.com/AlisaOmel/Tfh_network_discovery.

## Acknowledgments

J.D. was supported in part by NIAID DP2AI164325, NIAID R01AI170108, NIAID U01AI179514 and NHGRI U01HG012041. SP was supported in part by NIAID U19AI110495.

